# Inference and search on graph-structured spaces

**DOI:** 10.1101/2020.03.06.981399

**Authors:** Charley M. Wu, Eric Schulz, Samuel J Gershman

**Affiliations:** Harvard University; Max Planck Institute for Biological Cybernetics; Harvard University and the Center for Brains, Minds and Machines

**Keywords:** Function learning, Generalization, Inference, Graphs, Exploration-Exploitation

## Abstract

How do people learn functions on structured spaces? And how do they use this knowledge to guide their search for rewards in situations where the number of options is large? We study human behavior on structures with graph-correlated values and propose a Bayesian model of function learning to describe and predict their behavior. Across two experiments, one assessing function learning and one assessing the search for rewards, we find that our model captures human predictions and sampling behavior better than several alternatives, generates human-like learning curves, and also captures participants’ confidence judgements. Our results extend past models of human function learning and reward learning to more complex, graph-structured domains.

## Introduction

On September 15^th^, 1835, Charles Darwin and the crew of the HMS Beagle arrived in the Galapagos Islands. As part of a five-year journey to study plants and animals along the coast of South America, Darwin collected specimens of Galapagos finches, which would become an important keystone for his theory of evolution. Back in England, Darwin began to study the geographical distribution of the birds, particularly the relationship between their features and their habitat. He noticed that while finches on nearby islands had similar beaks (e.g., the vegetarian tree finches and the large insectivorous tree finches with their broad and stout beaks), finches on more distant islands were more dissimilar (e.g., the cactus ground finch with its long and spike-like beak). From these observations, Darwin concluded that these finches all originally derived from the same finch and then gradually adapted to the conditions of the Islands. Since nearby islands had similar conditions, finches on these islands had more similar beaks.

Darwin’s historical insight is an example of function learning, where a function represents a mapping from some input space to some output space. In Darwin’s case, the hypothesis was a function mapping a bird’s habitat to the characteristics of its beak (e.g., size). Function learning has traditionally been studied with continuous input spaces, but functions can also be defined over discrete input spaces such as graphs. While the geography of habitats can sometimes be described by a Cartesian coordinate system (latitude and longitude), the Galapagos is structured as a chain of islands, where the Euclidean distance within an island can be larger than the distance between islands. Since finches from the same island tend to be similar, the relevant metric for function learning may be topological rather than Euclidean distance, where the chain of islands can be described as a graph.

Function learning on graph-structured inputs spaces is not restricted to scientific epiphanies; it also applies ubiquitously to daily life. For example, the spread of disease, ideas, and cultural products from interpersonal contact can be understood as functions defined over social graphs. We can learn to predict which of our friends will like a piece of music after observing the music preferences of other friends in our social network. Similarly, as many parents of toddlers know, the appearance of a sickness in daycare is highly predictive of who will get sick next. Beyond social graphs, the flow of individuals in a transportation network and the distribution of food resources in patchy environments can likewise be described using graph-structured functions.

Despite the ubiquity of graph-structured functions, most studies of function learning (as we review below) have examined only continuous input spaces. In addition, reinforcement learning in discrete state spaces can also be interpreted as a form of graph-structured function learning, but relatively little work has examined patterns of generalization beyond very simple graph structures (e.g., Gershman & Niv, 2015; Wimmer, Daw, & Shohamy, 2012). Can similar computational principles of inference and search that describe human behavior in continuous spaces also apply to discrete, graph-structured spaces?

In this paper, we investigated how people learn graph-structured functions and use this knowledge to guide the search for rewards. In Experiment 1, we studied how people infer the values of nodes on complex graphs (corresponding to the number of passengers on a virtual subway map), where values were correlated by the connectivity structure, such that connected nodes had similar values. This is a discrete analogue of traditional function learning tasks on continuous input spaces, where we hypothesized that people would be able to make accurate predictions by taking into account the connectivity structure of the graph. We tested this hypothesis by analyzing performance and comparing different computational models in their ability to predict participants’ judgments and confidence ratings. In Experiment 2, we studied how people search for rewards on complex graphs, tantamount to a 64-armed bandit problem, where each arm of the bandit corresponded to a node on a graph and rewards were similarly correlated based on connectivity. Here, we hypothesized people would be able to leverage the structure of the environment to explore efficiently and rapidly acquire better rewards, using the same computational principles of function learning for inferring value. We tested this hypothesis through both behavioral analyses and computational modeling, where we compared models that differed in how they generalized about novel stimuli and in their exploration strategies.

Our results indicate that people learn and search for rewards consistent with a Bayesian model of function learning, implemented using Gaussian Process (GP) regression with a diffusion kernel. Our diffusion kernel GP model outperformed various alternatives in predicting inferences, uncertainty judgements, and when combined with an optimistic sampling strategy (upper confidence bound sampling), also performed best in predicting sampling decisions on a 64-armed bandit problem with graph-structured rewards. This model builds on past studies using Gaussian Process regression to describe human function learning on continuous spaces (Lucas, Griffiths, Williams, & Kalish, 2015; Schulz, Tenenbaum, Duvenaud, Speekenbrink, & Gershman, 2017), but using a prior over functions designed for discrete spaces (Kondor & Lafferty, 2002). Not only do we find strong empirical evidence for our model, but it also provides new theoretical connections to past research on human function learning, sample-efficient exploration, and classic theories of generalization and learning.

### Function learning in continuous spaces

Research on human function learning was originally pioneered by Carroll (1963), who studied how participants learned to predict the length of a line (output) based on the horizontal position of a “V” shaped marking (input). Unknown to participants, the relationship between the inputs and outputs were governed by either a positive linear, a quadratic, or a random function. Carroll’s (1963) study was motivated by the goal of showing that people could extrapolate functions to generate novel predictions about outcomes that had never before been observed. In contrast to classical theories of generalization (Shepard, Hovland, & Jenkins, 1961), Carroll’s work provided evidence for a mechanism of generalization that went beyond merely predicting the same outcome as that of the most similar previous experiences. Aside from showing that function learning was an important feature of human inference, Carroll (1963) also discovered that some functions, such as linear ones, were easier to learn than others, such as nonlinear ones. Subsequent studies of human function learning built on Carroll’s initial insight and further investigated which types of functions were more difficult to learn (Brehmer, 1974; Busemeyer, Byun, DeLosh, & McDaniel, 1997; Koh & Meyer, 1991), finding that linear functions with positive slopes are the most learnable, and that both nonlinear functions and linear functions with negative slopes are more difficult to learn.

A problem with many of these early studies was the inflexibility of their models. Likely inspired by timely advances in statistical methods of least-square estimation, they assumed that participants used a specific parametric model, for example linear regression, and then learned by optimizing the parameters to explain the data. Yet the parametric classes of function used in these studies were insufficiently flexible to account for human function learning. Instead of only adapting a specific class of functions to a particular set of observations, people seem to adapt the model itself when encountering novel data. Brehmer (1976) tried to explain some of these effects with a sequential hypothesis testing model of functional rule learning, according to which participants adapt the complexity of their model by performing sequential hypothesis tests and pivoting between parametric forms if necessary. However, this model still required a pre-determined set of parametric rules that could be compared, such that it is not able to explain the ability to learn almost any function given enough data. Thus, these earlier, *rule-based* models of human function learning could not easily explain the full range of human function learning abilities; more flexible models were needed.

To overcome the weaknesses of rule-based models of human function learning, several researcher proposed a novel class of *similarity-based* models of function learning. These models operated under the assumptions that similar input points will produce similar outputs and used neural networks to model behavior (McClelland, Rumelhart, & Group, 1986). These models could not only theoretically learn nearly any function, they were also able to capture the effect that linear functions are easier to learn than non-linear functions.

An important distinction in the literature on function learning (and machine learning more generally) is between interpolation (i.e., predictions for points nested between training examples) and extrapolation (i.e., predictions outside the convex hull of training inputs). Whereas similarity-based models can explain order-of-difficulty effects in interpolation tasks, they have trouble explaining how people extrapolate. Specifically, people tend to make linear predictions with a positive slope and an intercept of zero when extrapolating functions (Kwantes & Neal, 2006). This linearity bias holds true even when the underlying function is non-linear; for example, when trained on a quadratic function, average predictions fall between the true function and straight lines fit to the closest training points (Kalish, Lewandowsky, & Kruschke, 2004).

Since traditional similarity-based models of function learning could not easily explain these extrapolation patterns, the class of function learning models had to be extended even further. This led to the development of so-called *hybrid* models of function learning, which contain an associative learning process that acts on explicitly-represented rules. One such hybrid model is the Extrapolation-Association Model (DeLosh, Busemeyer, & McDaniel, 1997), which uses similarity-based interpolation, but extrapolates using a simple linear rule. The model effectively captured the human bias towards linearity, and could predict human extrapolations for a variety of functions, but without accounting for non-linear extrapolation (Bott & Heit, 2004).

More recently, another class of models was developed using *Gaussian Process* (GP; Rasmussen & Williams, 2006) regression to model function learning based on the principles of Bayesian inference. The GP framework describes a prior over functions, which given a set of observed data points, can be used to infer a posterior distribution over functions. Importantly, GP regression is a non-parametric model (Gershman & Blei, 2012; Schulz, Speekenbrink, & Krause, 2018), meaning that it adapts its complexity to the encountered data rather than assuming a fixed level of complexity. Griffiths, Lucas, Williams, and Kalish (2009) and Lucas et al. (2015) were the first to show that GP regression provides a rational model of human function learning, and that it replicates most of the observed empirical phenomena of human function learning. Importantly, GP regression performs posterior inference in a way that can be understood as both similarity-based (because the kernel provides a similarity metric between data points) and rule-based (because the kernel can be expressed as a linear weighted sum), providing a further unification of rule-based and similarity-based theories (Lucas et al., 2015).

### Using function learning to guide search

Learning a function is not only useful for making explicit generalizations about novel situations, but can also be used to guide adaptive behavior by leveraging functional structure to predict unobserved rewards in the environment. For example, in reinforcement learning tasks where options had inversely correlated rewards (Wimmer et al., 2012) or with rewards structured as a linear function (i.e., linearly increasing rewards from option 1 to option *N*; Schulz, Bhui, et al., 2019), participants were able to rapidly learn this structure and leverage it to facilitate better performance, even without having been explicitly told about the underlying structure.

In tasks with a large number of options, it becomes important to be able to learn efficiently, for instance by using features of the task to predict rewards (Farashahi, Rowe, Aslami, Lee, & Soltani, 2017; Radulescu, Niv, & Ballard, 2019). One approach is to learn an implicit value function mapping features onto rewards (Schulz, Konstantinidis, & Speekenbrink, 2017), which can be used to guide efficient exploration even in infinitely large problem spaces. Previous work has successfully used a GP model of function learning to predict human search behavior in a variety of both spatially and conceptually correlated reward environments (Schulz, Wu, Huys, Krause, & Speekenbrink, 2018; Wu, Schulz, Garvert, Meder, & Schuck, in press; Wu, Schulz, Speekenbrink, Nelson, & Meder, 2018), where the number of options vastly outnumbered the sampling horizon.

In transitioning from a pure function learning paradigm to a reward learning paradigm, the demands of the task change from pure information maximization to a balance between exploration and exploitation (Cohen, McClure, & Yu, 2007; Mehlhorn et al., 2015; Schulz & Gershman, 2019). Typically studied in multi-armed bandit tasks, the exploration-exploitation dilemma requires an agent to trade off between sampling novel options to acquire potentially useful information about the structure of rewards (exploration) with sampling options known to have high-value payoffs (exploitation). Not enough exploration, and the agent could get stuck in a local optima, while not enough exploitation and the agent never reaps the rewards they have discovered.

Since optimal solutions in such tasks are intractable for all but the most simplistic scenarios (Whittle, 1980), a variety of heuristic algorithms are commonly used. One such algorithm is upper confidence bound sampling, which adds an “uncertainty bonus” to each option’s value (Auer, 2002). Since this corresponds to a weighted sum of the expected reward and its uncertainty, this algorithm explicitly encodes the trade-off between exploration and exploitation. Although earlier studies produced mixed evidence for an uncertainty bonus in human decision making (Daw, O’doherty, Dayan, Seymour, & Dolan, 2006; Payzan-LeNestour & Bossaerts, 2011), many recent studies have shown that humans do engage in uncertainty-guided exploration (Gershman, 2018a, 2019; Knox, Otto, Stone, & Love, 2012; Speekenbrink & Konstantinidis, 2015; Wilson, Geana, White, Ludvig, & Cohen, 2014; Wu, Schulz, Speekenbrink, et al., 2018).

A key component for performing uncertainty-guided exploration is being able to estimate the uncertainty of one’s predictions. Since GP regression is a Bayesian model of function learning, uncertainty is quantified by the posterior distribution. In contrast, a model that makes only point estimates of expected reward does not have access to uncertainty-guided exploration. Instead, less efficient random exploration strategies must be used (e.g., softmax exploration). A combined model of GP regression with upper confidence sampling has proved to be an effective model in a wide number of contexts, describing how people explore different food options based on real world data (Schulz, Bhui, et al., 2019), predicting whether or not to people will try out novel options (Stojić, Schulz, Analytis, & Speekenbrink, 2020), and explaining developmental differences between how children and adults search for rewards (Meder, Wu, Schulz, & Ruggeri, 2020; Schulz, Wu, Ruggeri, & Meder, 2018).

### Function learning in graph-structured spaces

In the current work, we examine whether principles of function learning can be used to model human inference and search for rewards in graph-structured spaces (see Figure 1). Studying these environments greatly expands the scope of classical function learning models, and addresses an important gap in our understanding of how people reason about structured environments (e.g., social graphs or subway networks) that are ubiquitous in our daily lives.

**Figure 1.**
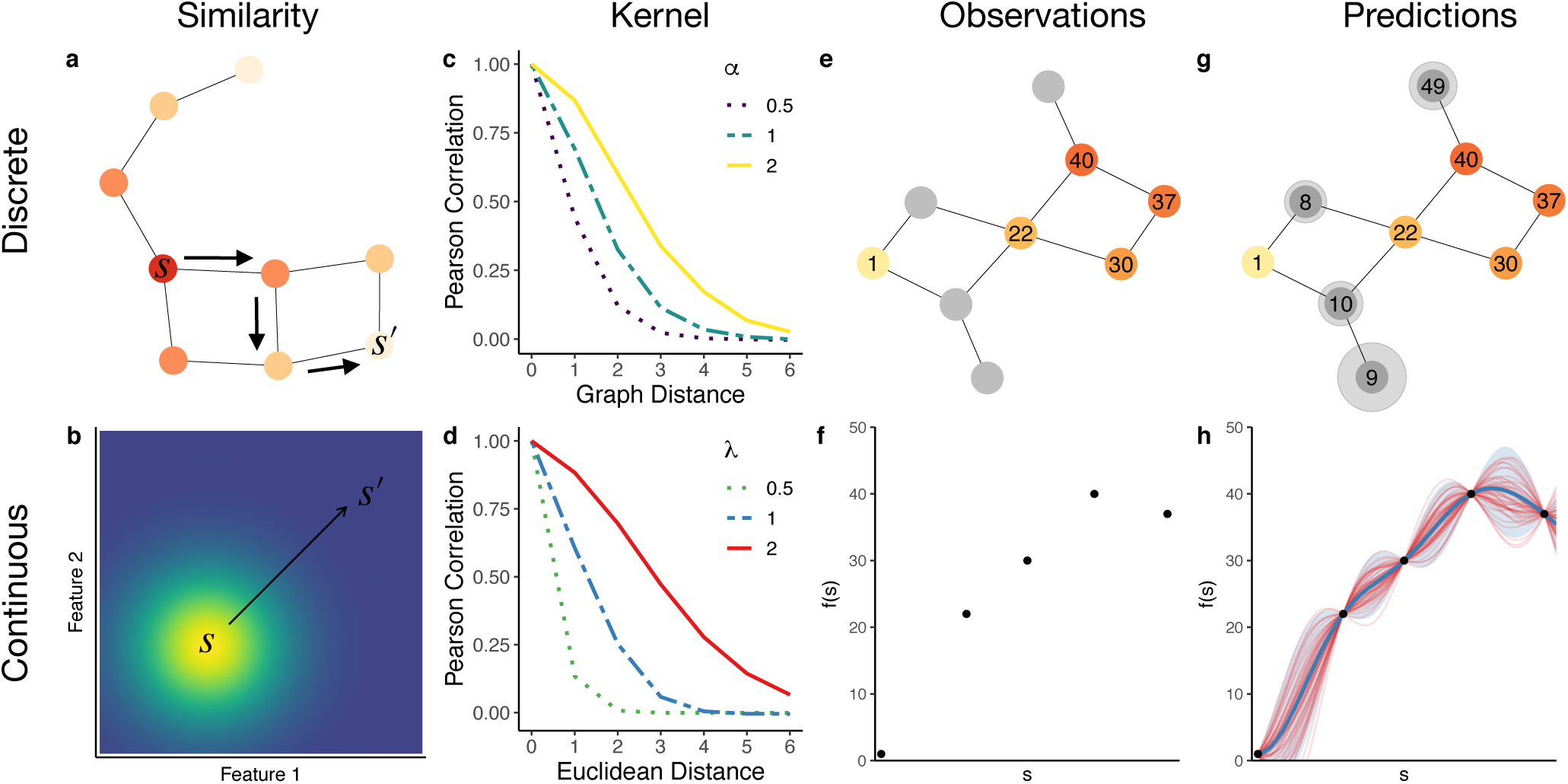
Function learning in discrete (top) and continuous (bottom) spaces. Moving from left to right, we illustrate how an appropriate measure of *similarity* encoded by a *kernel* function can be used infer a function, which when conditioned on previous *observations* makes Bayesian *predictions* about novel inputs. **a**) An example of a discrete graph structure, where nodes represent states, edges indicate transitions, and similarity is based on the connectivity between any two nodes *s* and *s*′. The color of the nodes indicates the covariance of the diffusion kernel *k*(*s, s*′) centered on node *s*, which decreases monotonically with graph distance. **b**) A continuous input space represents data as a set of feature values, where similarity is the inverse of (Euclidean) distance in feature space. Colors indicate the covariance of the radial basis function (RBF) kernel centered on input *s*, which decreases monotonically with Euclidean distance. **c**) The *diffusion kernel* assumes function values diffuse across the graph according to a random walk. The correlation between function values (normalized covariance) at any two nodes *s* and *s*′ decays monotonically as a function of graph distance. The diffusion parameter α governs the rate of decay. **d**) In the limiting case of an infinitely fine lattice graph, the diffusion kernel is equivalent to a *radial basis function* (RBF) kernel. The RBF kernel is commonly used in continuous domains, where covariance is a function of Euclidean distance between data points. The length-scale parameter λ controls the rate of this decay. **e**) Given some observations on a graph (colored nodes) or **f**) in a continuous input space (points), we use Gaussian process (GP) regression to make probabilistic predictions (**g-h**) about expected function values. **g**) In the graph example, the posterior mean is represented by numbers in the grey nodes, and the posterior uncertainty (variance) is represented by the size of the halo. **h**) In the continuous input space, the posterior mean is represented by the blue line, while the shaded blue ribbon represents the 95% confidence interval. Each of the red lines represents a randomly sampled function from the posterior.

In what follows, we will first introduce the GP regression framework, and then specialize it to the problem of function learning on graphs. The key mathematical tool that we employ is the diffusion kernel (Kondor & Lafferty, 2002), which offers one of the simplest ways to define priors over functions on graphs. We will show how the diffusion kernel naturally connects to past models of human function learning. We will then put this model to an empirical test, presenting two experiments studying how people make inferences and search for rewards on graph structures. In Experiment 1, participants were shown a series of artificially generated subway maps and asked to predict the number of passengers at unobserved stations. In Experiment 2, participants played a graph-structured multi-arm bandit task, where arms correspond to nodes in the graph, and the payoffs are correlated via the connectivity structure.

### Gaussian Process regression

A GP (Rasmussen & Williams, 2006) defines a distribution over functions **f** : 𝒮 → ℝ that map the input space *s* ∈ 𝒮 (e.g., nodes on a graph; Fig. 1a) to real-valued scalar outputs (e.g., rewards). Intuitively, for any finite set of inputs {*s*_1_, *s*_2_, …*s*_*N*_}, we can express the output of a function as a vector of finite length **f** = {*f*_1_, *f*_2_, …, *f*_*N*_}. Each function vector **f** can be modelled as a random draw from a multivariate normal distribution:

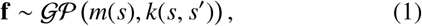

where *m*(*s*) = 𝔼[*f* (*s*)] is a mean function^1^ specifying the expected output of the function given input *s*, and *k*(*s, s*′) is the kernel function (see below) defining the covariance between outputs for a given input pair (*s, s*′).

We can think of each function as a potential hypothesis, relating each node *s* ∈ 𝒮 to some function value *f* (*s*), where the GP describes a distribution over functions and the kernel encodes inductive biases about how smoothly the function varies across the input space. Thus, the distributional nature of the GP captures uncertainty across different potential functions (Fig. 1). In the next section, we will define a kernel specialized for graph-structured functions.

We model a scenario in which an observer measures observations *y* = *f* (*s*)+*ϵ*, where *ϵ* ∼ *𝒩* (0, σ^2^) is Gaussian noise added to the output value. Given a data set of observations 𝒟 = **s, y** containing *N* input-output pairs (Fig. 1g-h), we can use a GP to compute the posterior predictive distribution *p*(*f* (*s*_∗_)| 𝒟) for any target state *s*_∗_ (e.g., an unobserved node; Fig. 1g). This posterior is a Gaussian distribution *p*(*f* (*s*_∗_)|𝒟) = *𝒩* (*m*_∗_, *v*_∗_), where the mean *m*_∗_ and variance *v*_∗_ are defined as:

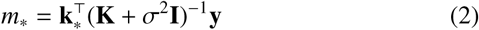

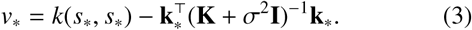

**K** is the *N* × *N* covariance matrix evaluated at each pair of observed inputs, and **k**_∗_ = [*k*(*s*_1_, *s*_∗_), …, *k*(*s*_*N*_, *s*_∗_)] is the covariance between each observed input and the target input *s*_∗_. Thus, as illustrated in Figure 1g, the GP posterior allows us to make Bayesian predictions about the expected output (*m*_∗_) and uncertainty (*v*_∗_) for any unobserved node *s*_∗_ on the graph.

As pointed out by Lucas et al. (2015), we can draw a connection between GP regression and similarity-based models of function learning. In particular, the posterior predictive mean (Eq. 2) can alternatively be expressed as a similarity weighted sum:

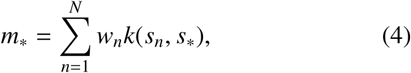

where each *s*_*n*_ is a previously observed input, and the weights are given by **w** = [**K** + σ^2^**I**]^−1^ **y**. Intuitively, this means that GP regression is equivalent to a linearly-weighted sum of similarities between the target input and the observed input (see Schulz, Speekenbrink, & Krause, 2018, for a tutorial).

### The diffusion kernel

We now introduce a kernel function that is specialized for graph-structured input spaces. A graph *G* = (*𝒮, ε*) consists of nodes *s* ∈ *𝒮* and edges *e*∈ *ε* (Fig. 1a). As a concrete example, a subway map describes a graph structure, where nodes correspond to stations and edges correspond to connections between stations. For now, we assume that all edges are undirected, so that probabilistic dependencies between any two adjacent nodes are symmetric.

The diffusion kernel (DF; Kondor & Lafferty, 2002) defines a similarity metric *k*(*s, s* ^′^) between any two nodes based on the matrix exponentiation^2^ of the graph Laplacian:

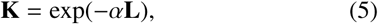

where the graph Laplacian **L** captures the transition structure of the graph based on the difference between the adjacency matrix **A** and degree **D**:

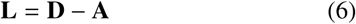

Each element *A*_*i j*_ of the adjacency matrix **A** is 1 when nodes *i* and *j* are connected, and 0 otherwise, while **D** is a diagonal matrix computed from the row sums of **A** and describe the number of connections of each node. Returning to our subway example, when there exists a route between stations *i* and *j*, then *A*_*i j*_ = 1 while *L*_*i j*_ = −1. In addition, for any station *i*, both *D*_*ii*_ and *L*_*ii*_ indicate the number of connected stations. The graph Laplacian can also describe graphs with weighted edges, where we can substitute the weighted adjacency matrix **W** for **A**, where each element *W*_*i j*_ describes the edge weight between nodes *i* and *j*, and the weighted degree of each node is expressed in the diagonals of **D**.

Intuitively, the graph Laplacian can be understood as a measure of the”flux” between nodes, for instance, the flow of passengers along different sections of a subway network. Flux between nodes *i* and *j* is not only influenced by whether they are connected, but is also affected by other connected nodes. For instance, if two train stations are connected to many other stations, then there is a relatively low probability that a randomly selected commuter will transit between them, compared to when the two stations have few alternative connections.

The diffusion kernel uses this intuition to define a similarity metric over discrete graph-structured spaces (Fig. 1a), by assuming that output values diffuse along the edges of a graph, similar to a heat diffusion process. The free parameter α → ∞ models the rate of diffusion, where α → 0 assumes complete independence between nodes, and αassumes all nodes are perfectly correlated. Thus, closely connected nodes are assumed to have similar output values, where the covariance between nodes decays monotonically as a function of graph distance (Fig. 1c).

### Connecting spatial and structured generalization

The GP framework allows us to relate similarity-based generalization on graphs to theories of generalization in continuous domains (Fig. 1 bottom row). Consider the case of an infinitely fine lattice graph (i.e., a grid-like graph with equal connections for every node and with the number of nodes and connections approaching infinity). Following Kondor and Lafferty (2002) and using the diffusion kernel defined by Eq. 5, this limit can be expressed as

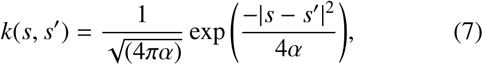

which is equivalent to the Radial Basis Function (RBF) kernel. The RBF kernel provides a similarity metric in continuous spaces based on Euclidean distance between data points (Fig. 1b), where similarity is the inverse of distance. In comparison, the diffusion kernel models similarity based on the dynamics of diffusion, where transitions are restricted by the graph structure. The RBF kernel can be understood as a special case of the diffusion kernel, when the environment is symmetric and transitions are unrestricted. The diffusion kernel is therefore able to offer a broader framework for modeling function learning and search, which subsumes past research on human behavior in spatial and conceptual input spaces (Wu et al., in press; Wu, Schulz, Speekenbrink, et al., 2018).

### Experiment 1: Subway prediction task

In our first experiment, participants were shown various graph structures described as subway maps (Fig. 2), and were asked to make predictions about unobserved nodes. For each prediction, participants also gave confidence judgments, which we use as an estimate of their (inverse) uncertainty. We used a GP parameterized with a diffusion kernel as a model of function learning in this task, which we compared to several alternative models.

**Figure 2.**
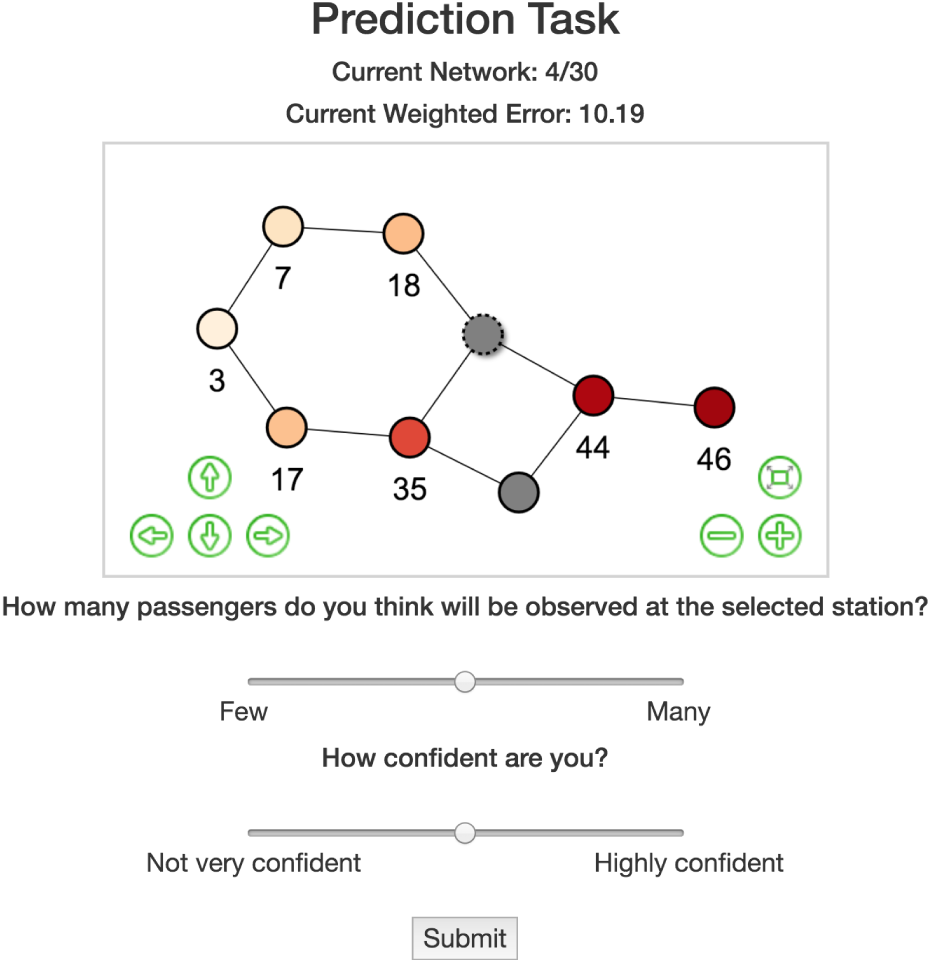
Screenshot from Experiment 1. Observed nodes (3, 5, or 7 depending on the information condition) are shown with a numerical value and a corresponding color aid. The target node is indicated by the dashed outline, and dynamically changed value/color as participants moved the top slider. Confidence judgments were used to compute a weighted error (i.e., more confident answers having a larger contribution), which was used to determine the performance-contingent bonus.

## Methods

### Participants

We recruited 100 participants (*M*_*age*_ = 32.7; *S D* = 8.4; 28 female) on Amazon Mechanical Turk (requiring a 95% approval rate and 100 previously completed HITs) to perform 30 rounds of a graph prediction task. The experiment was approved by the Harvard Institutional Review Board (IRB15-2048).

### Procedure

On each graph, numerical information was provided about the number of passengers at 3, 5, or 7 other stations (along with a color aid), from which participants were asked to predict the number of passengers at a target station (natural numbers from 0 to 50) and provide a confidence judgment (Likert scale from 1 to 11). The subway passenger cover story was used to provide intuitions about graph correlated functions, similar to our example from the introduction. The color aid was generated through a continuous, linear mapping (similar to Meder et al., 2020; Schulz, Wu, Ruggeri, & Meder, 2019; Wu, Schulz, Speekenbrink, et al., 2018), with both hue and brightness changing monotonically with value. Additionally, participants observed 10 fully revealed graphs to familiarize themselves with the task and completed a comprehension check before starting the task.

Participants were paid a base fee of $2.00 USD for participation with an additional performance contingent bonus of up to $3.00 USD. The bonus payment was based on the mean absolute judgement error weighted by confidence judgments: 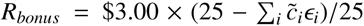 where 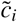 is the normalized confidence judgment 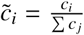 and *ϵ*_*i*_ is the absolute error for judgment *i*. On average, participants completed the task in 8.09 minutes (*S D* = 3.7) and earned $3.87 USD

All participants observed the same set of 40 graphs that were sampled without replacement for the 10 fully revealed examples in the familiarization phase and for the 30 graphs in the prediction task. We generated the set of 40 graphs by iteratively building 3 × 3 lattice graphs (also known as mesh or grid graphs), and then randomly pruning 2 out of the 12 edges. In order to generate the functions (i.e., number of passengers), we sampled a single function from a GP prior over the graph, where the diffusion parameter was set to α = 2.

### Modeling

We compared the predictive performance of the GP with two heuristic models that use a nearest-neighbors averaging rule (see below). Models were fit to each individual participant by using leave-one-round-out cross-validation to iteratively compute the maximum likelihood estimates on a test set, and then make out-of-sample predictions on the held-out round. We repeated this procedure for all rounds and compared the predictive performance (see Appendix B) over all held-out rounds.

The two heuristic strategies for function learning on graphs make predictions about the output values of a target state *s*_∗_ based on a simple nearest neighbors averaging rule. The *k-Nearest Neighbors* (kNN) strategy averages the values of the *k* nearest nodes (including all nodes with same shortest path distance as the *k*-th nearest), while the *d-Nearest Neighbors* (dNN) strategy averages the values of all nodes within path distance *d*. Both kNN and dNN default to a prediction of 25 when the set of neighbors are empty (i.e., the median value in the experiment).

Both the dNN and kNN heuristics approximate the local structure of a graph with the intuition that nearby nodes have similar output values. While they sometimes make the same predictions as the GP model while having lower computational demands, they fail to capture the full connectivity structure of the graph. Thus, they are unable to learn directional trends (e.g., increasing function values from one end of the graph to the other) or asymmetric influences (e.g., a central hub exerting relatively larger influence on sparsely connected neighbors). Additionally, they only make point-estimate predictions, and thus do not capture the underlying uncertainty of a prediction,which we use to model confidence judgments.

## Results and discussion

All code and data necessary to replicate the analyses in this manuscript are publicly available at https://github.com/charleywu/graphInference. Figure 3 shows the behavioral and model-based results of the experiment. We applied Bayesian mixed-effects regression to estimate the effect of the number of observed nodes on participant prediction errors, with participants as a random effect (see Table A1 for details). Participants made systematically lower errors in their predictions as the number of observations increased (bnumNodes = −0:60, 95% HPD: [−0:79; −0:41], BF10 =1.1 × 10^7^; Table A1; Fig. 3a). Repeating the same analysis but using participant confidence judgments as the dependent variable, we found that confidence increased with the number of available observations (*b*_numNodes_ = 0.23, 95% HPD: [0.17, 0.30], *BF*_10_ = 4.7 × 10^8^; Table A1; Fig. 3b). Finally, participants were also able to calibrate confidence judgments to the accuracy of their predictions, with higher confidence predictions having lower error (*b*_confidence_ = − 0.66, 95% HPD: [−0.83, −0.49], *BF*_10_ = 4.0 × 10^8^; Table A1; Fig. 3c). We found no effect of round number on prediction error (*b*_round_ = 0.01, 95% HPD: [0.02, −0.03], *BF*_10_ = 0.06), suggesting that the familiarization phase and cover story were sufficient for providing intuitions about graph correlated structures.

**Figure 3.**
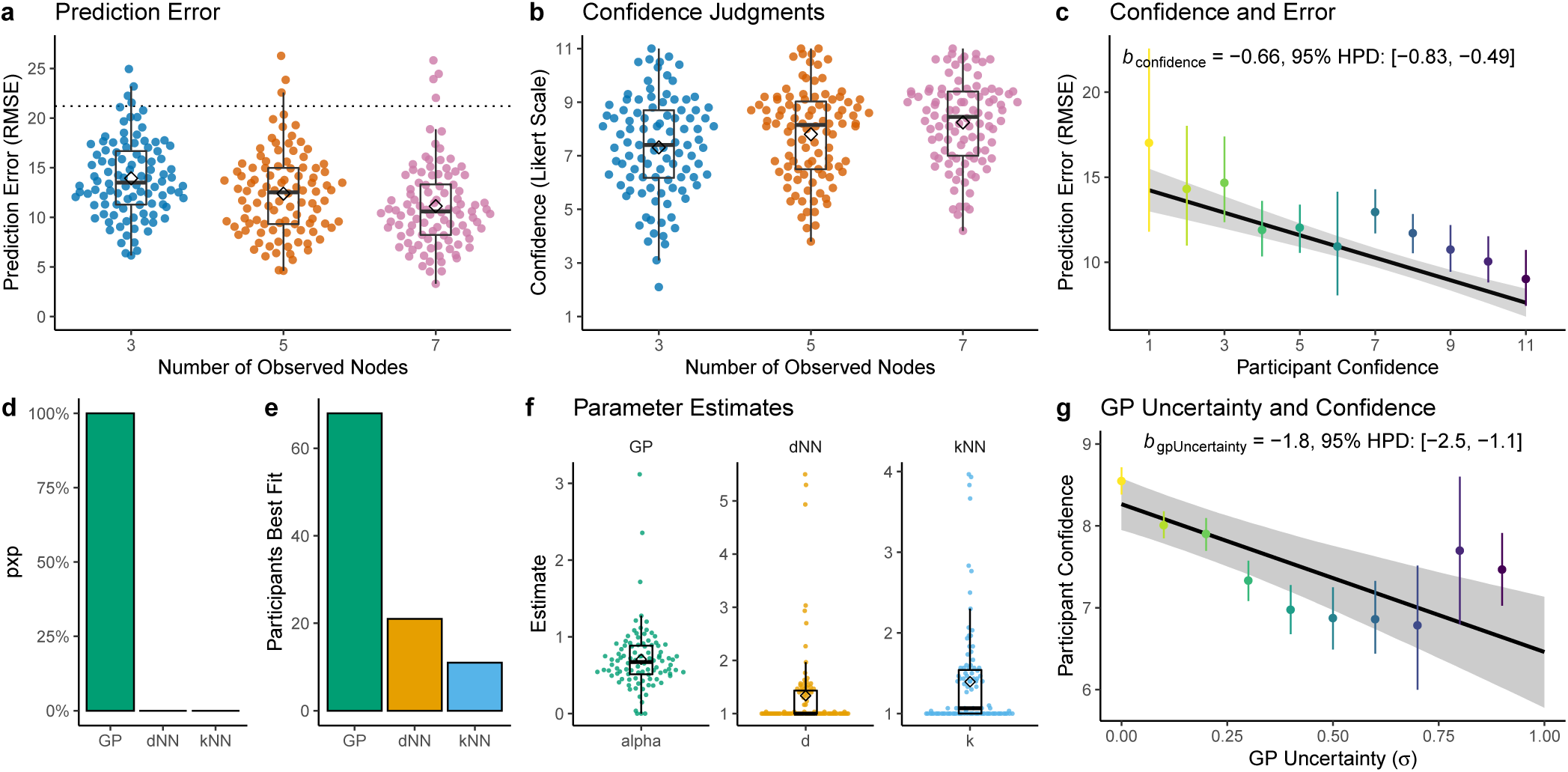
Experiment 1 results. **a-b**) Participant judgment errors and confidence estimates. Each dot is a single participant (averaged over each number of observed nodes), with Tukey box plots and diamonds indicating group means. The dotted line in **a**) is a random baseline (simulated by uniformly sampling judgments from 0-50). **c**) Judgment error and confidence. Each colored dot is the aggregate mean with error bars indicating the 95% CI. The black line is the group-level effect of a Bayesian mixed model (Table A1), indicating the posterior mean and 95% CI (ribbon). **d**) Hierarchical Bayesian model comparison between the Gaussian process (GP) with diffusion kernel, d-nearest neighbors (dNN), and k-nearest neighbors (kNN). The bars indicate the protected exceedence probability (pxp) as an estimate of the posterior probability of a given model being the most frequent in the population (corrected for chance). **e**) The number of participants best fit by each model. **f**) Parameter estimates, where each dot is the mean cross-validated estimate for each participant, with Tukey box plots and diamonds indicating group means. **g**) The inverse relationship between GP uncertainty estimates (σ) and participant confidence judgments (Likert scale), where the colored dot and error bars indicate (respectively) the aggregate mean ± 95% CI computed at 10 equally spaced intervals along the x-axis. The black line is the fixed-effect of a Bayesian mixed model (Table A1), with the ribbon indicating the 95% CI.

Figure 3d shows the model comparison results. We evaluated the relative performance of models using the protected exceedence probability (pxp), as a Bayesian estimate of the probability that a particular model is more frequent in the population than all the other models under consideration, corrected for chance (see Appendix A; Rigoux, Stephan, Friston, & Daunizeau, 2014; Stephan, Penny, Daunizeau, Moran, & Friston, 2009). The GP with diffusion kernel was overwhelmingly the best model, with *pxp*(*GP*) ≈ 1. Overall, 68 out of 100 participants were best predicted by the GP, 21 by the dNN, and 11 by the kNN (Fig. 3e; see Fig. B1 for additional comparisons between model predictions and participant judgments).

Figure 3f shows individual parameter estimates of each model. The estimated diffusion parameter α was substantially lower than the ground truth of α = 2 (*t*(99) = −31.3, *p* < .001, *d* = 3.1, *BF*_10_ = 4.4 × 10^29^)^3^, replicating previous findings that have shown undergeneralization to be a prominent feature of human behavior (Wu, Schulz, Speekenbrink, et al., 2018). Estimates for *d* and *k* were highly clustered around the lower limit of 1, suggesting that averaging over larger portions of the graph were not consistent with participant predictions.

Lastly, an advantage of the GP is that it produces Bayesian uncertainty estimates for each prediction. While the dNN and kNN models make no predictions about confidence, the GP’s uncertainty estimates correspond to participant confidence judgments, which we validated using a Bayesian mixed model regressing the uncertainty estimates of the GP onto participant confidence judgments (*b*_*gpUncertainty*_ = −1.8, 95% HPD: [−2.5, −1.1], *BF*_10_ = 1.2 × 10^5^; Table A1, Fig. 3g).

The results of this experiment demonstrate that a GP with a diffusion kernel can successfully model human function learning on graphs, in particular the empirical pattern of predictions and confidence ratings. Our model extends existing theories of human function learning in continuous spaces, where the RBF kernel (commonly used in continuous domains) can be seen as a special limiting case of the diffusion kernel.

### Experiment 2: Graph bandit

In our next experiment, we tested the suitability of the diffusion kernel as a model of search, using a multi-armed bandit task with structured rewards (see also Wu, Schulz, Speekenbrink, et al., 2018). In particular, extending our previous work on spatially and conceptually correlated multiarmed bandits (Wu et al., in press; Wu, Schulz, Speeken-brink, et al., 2018), we constructed a task where rewards were defined by the connectivity structure of a graph (Fig. 4). In this task, participants searched for rewards by clicking nodes on a graph. As in Experiment 1, the output values (rewards) were generated by a function drawn from a GP with a diffusion kernel. This induced a graph-correlated reward structure, allowing for similarity-based generalization to aid in search, but where similarity was defined based on connectivity rather than perceptual features or Euclidean distances between options as in our previous work.

**Figure 4.**
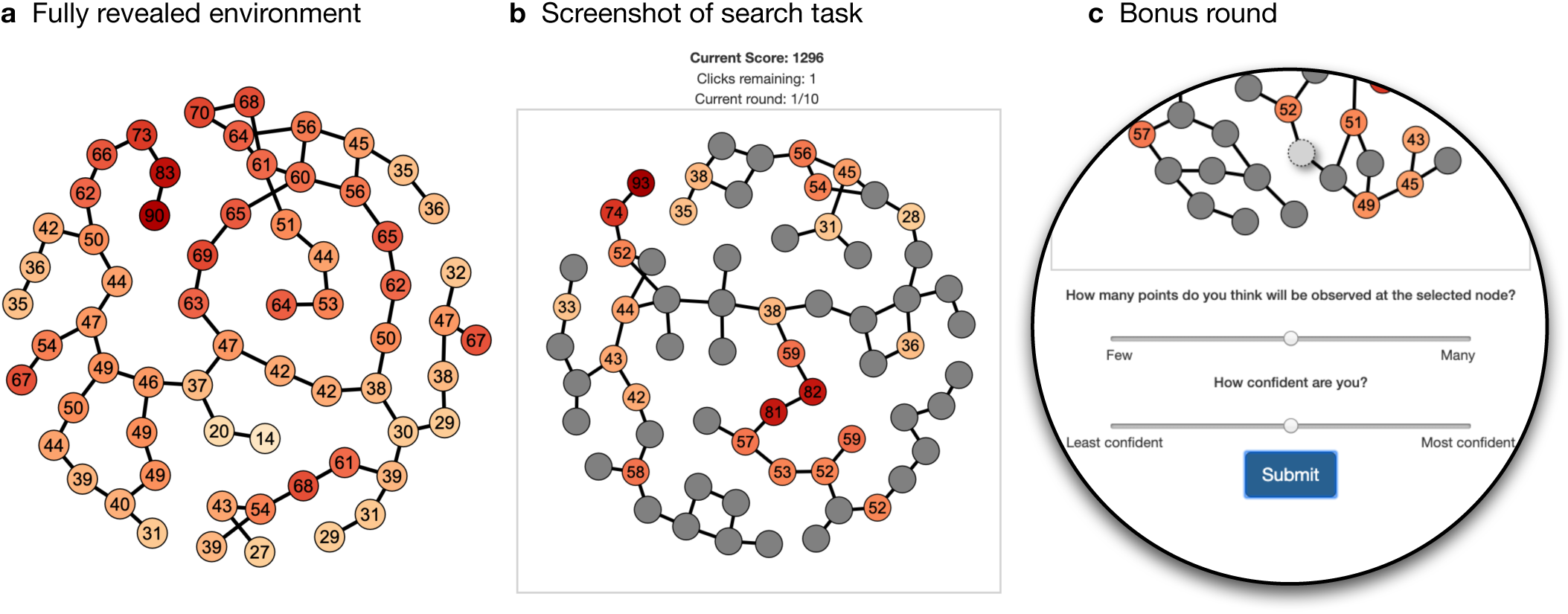
Experiment 2 screenshots. **a**) Four fully revealed environments were shown to participants prior to beginning the task. **b**) During the task participants were instructed to click nodes to earn as much reward as possible. Clicked nodes displayed the numeric value of the earned reward and a color guide (darker colors indicate higher rewards). **c**) Zoomed in screenshot of the bonus round, which activated after the 20th trial on the last round. 10 unclicked nodes were uniformly sampled and participants were sequentially asked to make judgments about expected rewards and report their confidence rating. The expected reward slider was mapped to the selected node, such that the color and numerical value dynamically changed as the slider was moved.

## Methods

### Participants

We recruited 100 participants on Amazon Mechanical Turk (requiring 95% approval rate and 100 previously completed HITs). Two participants were excluded because of missing data, making the total sample size *N* = 98 (*M*_*age*_ = 34.3; *S D* = 8.7; 32 female). Participants were paid $2.00 for completing the task and earned an additional performance contingent bonus of up to $3.00. Overall, the task took 7.2 ± 3.3 minutes and participants earned $4.32 ± $0.24 USD. The experiment was approved by the Harvard Institutional Review Board (IRB15-2048).

### Procedure

Participants were instructed to earn as many points as possible by clicking on the nodes of a graph. Each node represented a reward generating arm of the bandit, where connected nodes yielded similar rewards, such that across the whole graph the expected rewards were defined by a graph-correlated structure (see Fig. 4a). Along with the instructions indicating the correlated structure of rewards, participants were shown four fully revealed graphs to familiarize them with the reward structure and had to correctly answer three comprehension questions before starting the task.

After completing the comprehension questions, participants performed a search task over 10 rounds, each corresponding to a different randomly sampled graph structure.

In each task, participants were initially shown a single randomly revealed node, and had 25 clicks to either explore unrevealed nodes or to reclick previously observed nodes, where each observation included normally distributed noise *ϵ* ∼ *𝒩* (0, 1). Each clicked node displayed the numerical value (most recent observation if selected multiple times) and a color aid, where darker colors corresponded monotonically to larger rewards (Fig. 4b). After finishing each round, participants were informed about their performance as a percentage of the best possible score (compared to selecting the global optimum each trial). The final performance bonus (up to $3.00) was also calculated based on this percentage, averaged over all rounds.

In total, we generated 40 different graphs by building 8×8 lattice graphs and then randomly pruning 40% of the edges, with the constraint that the resulting graph be comprised of a single connected component. We then sampled a single reward function for each graph from a GP prior, parameterized by a diffusion kernel fit on the graph (with α = 2). The layout for each graph was pre-generated using the Fruchterman-Reingold (1991) force-directed graph placement algorithm, such that a single canonical layout for each graph was observed by all participants. For each participant, we sampled (without replacement) from the same set of 40 pre-generated graphs to build the set of 4 fully revealed graphs shown in the instructions and the 10 graphs used in the main experiment.

Prior to beginning the very last round, participants were informed that it was a “bonus round”. The goal of acquiring as many points as possible remained the same, but after 20 clicks, participants were shown a series of 10 unrevealed nodes and asked to make judgments about the expected reward and their confidence (Fig. 4c). After all 10 judgments were completed, participants were forced to choose one of the 10 options, and then the task was completed as normal. Behavioral and modeling results exclude the bonus round, except for the analyses of the judgment data.

### Modeling

In order to understand how participants search for rewards, we used computational modeling to make predictions about choices in the bandit task and the judgments from the bonus round. Models were fit to the bandit data (omitting the bonus round) using leave-one-round-out cross validation, where we iteratively held out a single round as a test set, and computed the maximum likelihood estimate on the remaining rounds as the training set. We compared models using the summed out-of-sample prediction accuracy on the held-out rounds. Altogether, we compared four different models corresponding to different strategies for generalization and exploration (see below).

Each model computes a value for each option *q*(*s*), which is then transformed into a probability distribution using a softmax choice rule:

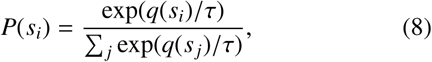

where the temperature parameter *τ* is a free parameter controlling the level of random exploration. In addition, all models also use a stickiness parameter *ω* that adds a bonus onto the value of the most recently chosen option. This is a common feature of reinforcement learning models (Christakou et al., 2013; Gershman, Pesaran, & Daw, 2009) and particularly in multi-armed bandit tasks (Schulz, Bhui, et al., 2019), which we include here to account for repeat clicks.

The GP model uses the diffusion kernel (Eq. 5) to make predictive generalizations about reward, where we fit α as a free parameter defining the extent to which generalizations diffuse along the graph structure. For each node *s*, the GP produces normally distributed predictions that can be summarized in terms of an expected value *m*(*s*) (Eq. 2) and the underlying uncertainty *v*(*s*) (Eq. 3). In order to model how participants balance between exploiting high value rewards and exploring highly uncertain options, we use upper confidence bound (UCB) sampling (Auer, 2002) to produce a valuation of each node:

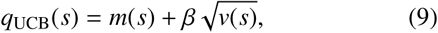

where the exploration bonus *β* is a free parameter that governs the level of exploration directed towards highly uncertain options. Higher values of *β* correspond to more exploratory behavior, which is directed towards nodes with the highest estimated level of uncertainty.

The *Bayesian mean tracker* (BMT) is a prototypical reinforcement learning model that can be interpreted as a Bayesian variant of the traditional Rescorla-Wagner (1972) model (Gershman, 2015). Like the GP, the BMT assumes a normally distributed prior over rewards *𝒩* (*m*_*j*,0_, *v* _*j*,0_). However, unlike the GP, the BMT assumes independent priors for each option *j*, where we set the prior mean to the median value of payoffs *m*_*j*,0_ = 50 and the prior variance to *v* _*j*,0_ = 500.

Given some observations 𝒟_*t*−1_ = {**X**_*t*−1_, **y**_*t*−1_} of rewards **y**_*t*−1_ for inputs **X**_*t*−1_, the BMT learns the rewards of each option by computing independent posterior distributions for the expected reward µ _*j*_ for each option *j*

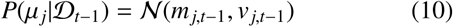

Thus, the BMT does not generalize but rather updates each posterior mean *m*_*j,t*_ and variance *v* _*j,t*_ independently:

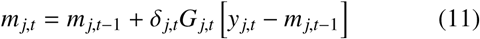

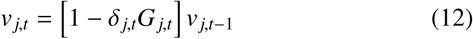

where the Kronecker delta δ _*j,t*_ = 1 if option *j* was chosen on trial *t*, and 0 otherwise, meaning the posteriors for the unchosen options are unchanged. Intuitively, the estimated mean of the chosen option *m*_*j,t*_ is updated based on the difference between the observed value *y*_*t*_ and the predicted mean from the previous time point *m*_*j,t*−1_ (i.e., prediction error). This update is scaled by a learning rate known as the Kalman gain *G*_*j,t*_:

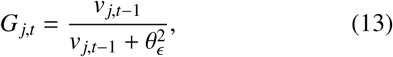

defined as a ratio of the prior variance *v* _*j,t*−1_ and the error variance 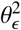. While the estimated mean is updated based on prediction error, the estimated variance for the chosen option *v* _*j,t*_ is reduced by a factor of 1 − *G*_*j,t*_, which is in the range [0, 1]. The error variance 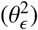 is treated as a free parameter and can be interpreted as inverse sensitivity. Smaller values of 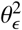 therefore result in more substantial updates to the mean *m*_*j,t*_, and larger reductions of uncertainty *v* _*j,t*_.

Like the GP, the BMT also used UCB sampling, along with stickiness and a softmax choice rule. Unlike the GP, the BMT does not generalize, and thus defaults to the prior mean and variance for any unobserved options. For this reason, we did not consider the BMT as a candidate model for Exp. 1. Nevertheless, it provided a sensible benchmark in the graph bandit task, since it is an optimal model for learning independent reward distributions through experience, and can support both directed and random exploration algorithms.

In addition to reinforcement learning models, we also considered both the dNN and kNN heuristics from Experiment 1. Predictions about expected reward were computed using the respective nearest neighbor averaging rule, where *d* and *k* were estimated as free parameters. For predictions where no observed nodes satisfied the averaging rule (i.e., all observations were too far away), we defaulted to an expected value of *m*(*s*) = 50 (median over all environments). In contrast to the GP and BMT models, these models make only point estimates about reward, and thus precluded UCB sampling. Instead, choice probabilities were calculated using only softmax choice rule on expected reward and with estimated stickiness weights.

## Results and discussion

Participants performed well in the task, achieving higher rewards over successive trials (*r* = .93, *p* < .001, *BF*_10_ = 4.5 × 10^7^; Fig. 5a) and decisively outperforming a random baseline (*t*(97) = 29.6, *p* < .001, *d* = 3.0, *BF*_10_ = 7.2× 10^46^). There was no substantial evidence for an influence of round number on performance (*r* = .49, *p* = .182, *BF* = 1.1), indicating that the fully revealed environments in the instructions (Fig. 4a) and comprehension questions were sufficient for conveying the goal of the task and the underlying covariance structure of rewards.

**Figure 5.**
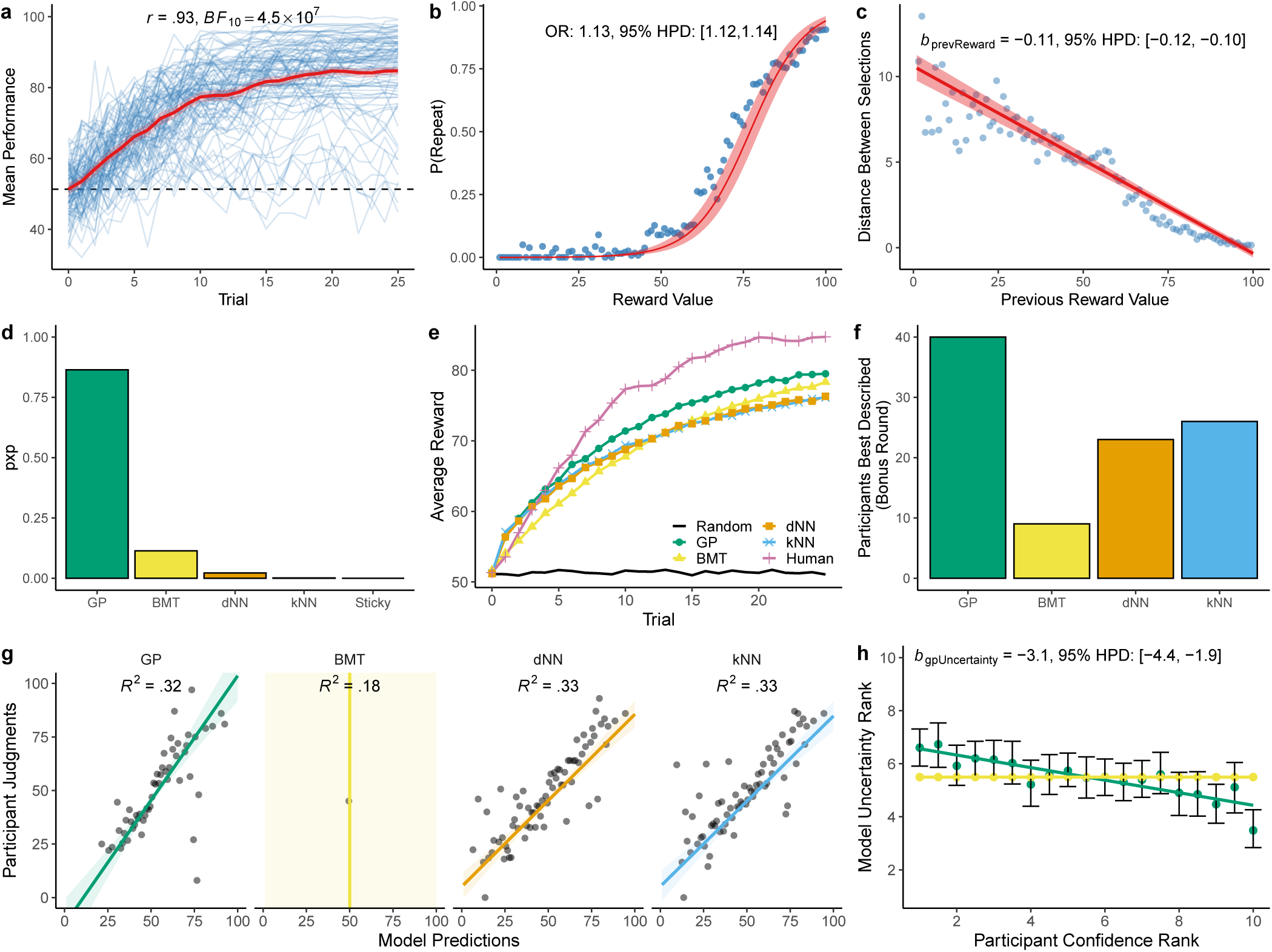
Experiment 2 results. **a**) Mean performance over trials, where each blue line is a single participant and the red line is the group mean (± 95% CI). The dashed line provides a comparison to a random baseline. **b**) The probability of repeating a selection as a function of the reward value. Each dot is the aggregate mean calculated at intervals of 1, while the red line is the group-effect of a Bayesian mixed effects logistic regression (Table A2), with the ribbon indicating the 95% CI. **c**) The relationship between reward value and the graph distance (shortest path) to the subsequent selection. Each dot is an aggregate mean, while the red line indicates the group-level effect of a Bayesian mixed model (Table A2). **d**) Model comparison based on out-of-sample predictive accuracy. We compare the Gaussian process (GP) model, the Bayesian mean tracker (BMT), *d*-Nearest Neighbors (dNN), *k*-Nearest Neighbors (kNN), and a stickiness-only (Sticky) model. The Y-axis shows the protected exceedance probability (pxp) describing the prevalence of each model in the population (corrected for chance). **e**) Simulated learning curves by sampling (with replacement) from participant parameter estimates (10k replications), where the black line shows a random baseline and the pink line shows mean participant performance. **f**) Participants best described on the bonus round data, where we used parameter estimates from the bandit task to make predictions of participant judgments about unobserved rewards on the bonus round and compared RMSE. **g**) The correspondence between each bonus round judgment and model predictions, where each dot is the aggregate mean calculated in intervals of 1, and each line is the group-level effect of a Bayesian mixed model (with ribbon indicating 95% CI). The Bayesian *R*^2^ of each mixed model is reported (see Table A3 for details). **h**) Only the GP and BMT make uncertainty estimates. Here we show the correspondence between rank ordered (per participant) confidence judgments and model uncertainty estimates. Dots indicate means with error bars showing 95% CI, and colored lines represent a linear regression. The regression coefficient corresponds to a Bayesian mixed model fit to the raw, untransformed data (see Table A3).

Participants adapted their search behavior as a function of reward value: higher rewards predicted a higher probability of making a repeat selection (Bayesian mixed model: Odds ratio = 1.13, 95% HPD: [1.12,1.14], *BF*_10_ = 3.2 × 10^40^; Table A2; Fig. 5b). We also found that higher rewards predicted shorter path distances to the subsequent selection (*b*_*prevReward*_ = − 0.11, 95% HPD: [−0.12, −0.10], *BF*_10_ = 2.6× 10^43^; Fig. 3c). Thus, participants searched locally when finding high rewards, and explored further way upon finding poor rewards (see Appendix C for analyses on connectivity structure and sampling patterns). This provides early evidence that participants used generalization to guide their search for rewards, since they systematically adapted their search distance as a function of reward value, thereby avoiding regions with poor rewards and searching more locally in richer areas.

Overall, the GP was the most predictive model (Fig. 5d) with an estimated prevalence of *pxp*(*GP*) = .86, with the other models having *pxp*(*BMT*) = .11, *pxp*(*dNN*) = .02, and *pxp*(*kNN*) = .001. As a benchmark, we also fit a null model that made the same prediction for every node, which combined with stickiness and the softmax choice rule, was worse than all other models *pxp*(*sticky*) < .001. At the individual level, 34 out of 98 participants were best fit by the GP, 27 by the BMT, 20 by the dNN, and 17 by the dNN. Participants with higher performance on the bandit task were better predicted by the GP model (*r* = −.83, *p* < .001, *BF* = 8.8 × 10^21^) and also tended to be more diagnostic between the GP and BMT models (see Fig. D3), in favor of the GP.

We simulated the behavior of each model by sampling (10k samples with replacement) from the set of participant parameter estimates and computing the average learning curves (Fig. 5e). Although all models performed below the human curves, the GP achieved the closest levels of performance, with the BMT performing next best (see Appendix D for a detailed analysis of parameter estimates from each model).

To provide additional support for our modeling results, we also predicted participant judgments in the bonus round using participant parameter estimates from the bandit task. Since model parameters were estimated through cross-validation on all rounds except the bonus round, we used each participant’s median parameter estimates (over rounds 1 to 9) to make out-of-task predictions about the bonus round judgments. The GP model best predicted the largest number of participants (Fig. 5f) and had the lowest prediction error on average (comparing RMSE), although there was no difference in comparison to the dNN (*t*(97) = −0.1, *p* = .897, *d* = 0.01, *BF* = .11), which had the second lowest prediction error. Looking more closely at the individual correspondence between participant judgments and model predictions (Fig. 5g), we fit separate Bayesian mixed effects regression for each model, predicting participant judgments based on model predictions (Table A3). Overall, the fits of these models for the GP, dNN, and kNN were highly similar.

While there was mixed evidence for which model best predicted judgments of expected reward, we next looked at how predictions of uncertainty corresponded to participant confidence ratings. Here, we interpreted confidence to be the inverse of uncertainty. In this analysis, we considered only the GP and BMT models, since no other models could generate uncertainty estimates. Figure 5h shows a comparison between the (per participant) rank-ordered confidence ratings and rank-ordered uncertainty estimates of the models. While the BMT estimated the same level of uncertainty for all unobserved nodes (making correlations undefined), we found that the GP uncertainty estimates corresponded well with participant confidence ratings. In order to test this relationship by accounting for individual differences in subjective ratings of confidence, we fit a mixed effects model to predict the raw confidence judgment (Likert scale 1-11) using the GP uncertainty estimate as a fixed effect and participant as a random effect (Table A3). The results showed a strong correspondence between lower confidence ratings and higher GP uncertainty estimates (*b*_*gpUncertainty*_ = −3.1, 95% HPD: [−4.4, −1.9], *BF*_10_ = 4.5 × 10^5^).

To summarize, Experiment 2 showed that participants leverage their functional knowledge over graph-structured reward environments to guide sampling decisions. Participants searched for rewards locally and found highly rewarding nodes much faster than would be expected under the assumption of independent options. We again found support for a GP model of function learning, augmented with a probabilistic action policy including both directed and undirected exploration. The GP provided the best predictive accuracy of choices, produced similar learning curves to human performance, and accurately predicted judgments about expected reward and confidence.

Nevertheless, we also found that the two nearest neighbor models closely matched the GP in terms of predicting choices and participants’ judgements of unobserved nodes. However, only the GP model can generate predictions of uncertainty, which we found to match well with participants’ confidence judgments. Given that we also found lower levels of α compared to the ground truth, participants likely generalized only very locally. Yet they still tracked their uncertainty about different options, and used that uncertainty to guide their exploration and to rate the confidence of their own predictions. The ability to model these characteristics of human behavior is what makes the GP model a superior model of human behavior in our task.

### General discussion

We studied how people learn and exploit graph-structured functions in two experiments. In Experiment 1, we studied how people make predictions about the values of unobserved nodes on a graph and estimated their level of confidence. In Experiment 2, we studied how people searched for rewards in a multi-armed bandit task with graph-correlated rewards. In both experiments, we found that participants made inferences, rated confidence, and navigated the exploration-exploitation dilemma consistent with a Bayesian model of human function learning. This model is implemented using GP regression, which has previously been shown to accurately describe function learning in continuous domains. Here, we replaced the RBF kernel commonly used in continuous domains with a diffusion kernel, where connectivity rather than feature similarity defines relationships in structured environments. The diffusion kernel in turn contains the RBF kernel as a special case, where any Cartesian feature space is equivalent to an infinitely fine undirected lattice graph. Thus, our model expands upon past research on human function learning to richer, graph-structured domains.

Our work also relates directly to classical work on human generalization (Shepard, 1987). Just as in Shepard’s original theory, the diffusion kernel defines a distance-dependent similarity measure which assumes that the similarity between nodes decays with their (graph) distance. Similar mechanisms have permeated theories of category learning, where participants learn about a stimulus class given its features (Kruschke, 1992; Love, Medin, & Gureckis, 2004; Medin & Schaffer, 1978; Nosofsky, 1984). Indeed, similar models have also been used to explain human decision making. For example, Gureckis and Love (2009) showed that participants can use the covariance between changing payoffs and the systematic change of a state cue to generalize experiences from known to novel states, and that a linear network learning using similarities between features matched well with participants’ behavior. By further combining the diffusion kernel with traditional models of generalization and category learning, we hope to pave the way towards a truly unifying theory of human generalization (Shepard, 1987; Tenenbaum & Griffiths, 2001; Wu, Schulz, Garvert, Meder, & Schuck, 2018).

Most directly related to our work is the theory of property induction developed by Kemp and Tenenbaum (2009), who showed how different assumptions about graph-structured functions can lead to different patterns of generalization, consistent with human data. For example, assumptions about genetic transmission through a taxonomic tree license different patterns of generalization compared to assumptions of disease transmission through a food chain. Whereas Kemp and Tenenbaum studied binary property induction, we have focused on real-valued properties in this paper. We have also gone beyond induction to study the role of structured function learning in decision making.

Recent work in reinforcement learning has also developed models related to the diffusion kernel. In particular, cumulative rewards can be estimated efficiently using the successor representation (SR), which represents states of the environment in terms of the expected future occupancy of other states (Dayan, 1993; Gershman, 2018b; Stachenfeld, Botvinick, & Gershman, 2017). For example, a particular subway station would be represented by a vector encoding the expected future occupancies of other stations in the network. When an agent follows a random walk in state space (approximating a diffusion process), the SR is equivalent to the inverse graph Laplacian. Thus, while it does not make probabilistic predictions about cumulative reward values (but see Geerts, Stachenfeld, & Burgess, 2019), the SR is able to generalize based on the diffusion of cumulative rewards in a graph-structured state space.

One limitation of our current model implementation is that we assumed the graph structure to be known *a priori*. While this may be a reasonable assumption in problems such as navigating a subway network, where maps are readily available, this is not always the case. One promising avenue for future research is to combine our model with other approaches that learn the underlying structure from experience. The Bayesian structure learning framework proposed by Kemp and Tenenbaum (2008) learns a “conceptual universe” of different graphs. The Bayesian model assigns a score to each candidate graph based on its prior probability and its likelihood. The prior is specified by a generative model for graphs (which can generate grids, trees, and chains, among other graphs) that favors simple, regular graphs over complex ones. The likelihood is based on the match between the observed data and the graph structure, under the assumption that the feature values of the data vary smoothly over the graph. In particular, the features are assumed to be multivariate Gaussian distributed with a covariance function defined by a variant of the regularized Laplacian kernel (Smola & Kondor, 2003; Zhu, Lafferty, & Ghahramani, 2003), which is closely related to the diffusion kernel used here.

Another limitation of our current study is that several variants of a simple nearest-neighbor averaging rule were also surprisingly effective heuristics to capture human behavior in our tasks. Both the dNN and kNN can be understood as binarized simplification of the similarity metric used by the GP. While the GP predicts expected rewards using a similarity-weighted sum of previous observations (Eq. 4), the dNN and kNN use either a distance or count-based threshold of similarity, such that nodes are either similar if considered a neighbor, or dissimilar otherwise. Similar nodes are then averaged, equivalent to an equal-weight regression model (Lichtenberg & Simsek, 2016; Wesman & Bennett, 1959). Although these heuristics are able to efficiently capture many aspects of judgments and choices, our results also show that human behavior is sensitive to the uncertainties about their own predictions, using them to rate their own confidence and to preferentially explore more uncertain options when searching for rewards. This aspect could only be captured by the GP model, which can generate Bayesian uncertainties about its own predictions. Future studies could try to create further heuristic models that also calculate uncertainties of different nodes, for example by using Bayesian versions of the nearest neighbor algorithms (Behmo, Marcombes, Dalalyan, & Prinet, 2010).

Currently, we have only focused on using the diffusion kernel for modeling smooth functions on graph structures. However, the real world contains mixes of continuous and discrete structures, such as in our example of Darwin’s finches. How could we model these more complex mixtures of structures? Participants’ ability to learn more complex yet highly structured functions in a continuous domain have been explained by using compositional kernels (Schulz, Tenenbaum, et al., 2017). Compositional kernels learn about functions through combining different structures, starting from simple building blocks that can be composed. Thus, one avenue for future research could be to model human function learning using kernel that compose together these mixtures of graph and continuous structures.

Finally, even under the assumption that participants know the full graph structure, our model additionally assumes that they have direct access to the value of each node during inference. Future studies could try to further enrich our tasks to scenarios in which participants have to plan moves over the graph, or to require that participant remember previously observed outputs. While studying how people plan on known graphs would connect our work further to past research on hierachical planning in human reinforcement learning (Balaguer, Spiers, Hassabis, & Summerfield, 2016; Tomov, Yagati, Kumar, Yang, & Gershman, 2018), adding a forgetting component to our model and task would connect it to memory-based models of learning (Bornstein & Norman, 2017; Collins & Frank, 2012) and decision making (Bhui, 2018; Stewart, Chater, & Brown, 2006).

In summary, our behavioral results and proposed model considerably expand past studies of human function learning to graph-structured domains, and emphasize the importance of function learning and uncertainty-guidance to explain human behavior in such domains.

## Appendix A Statistics and regression models

### Comparisons

We report both frequentist and Bayesian statistics. Frequentist tests are reported as Student’s *t*-tests (specified as either paired or independent). The Bayesian variant uses the default two-sided Bayesian *t*-test for either independent or dependent samples, where both use a Jeffreys-Zellner-Siow prior with its scale set to 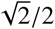 (Rouder, Speckman, Sun, Morey, & Iverson, 2009). Each test is accompanied by a Bayes factors (*BF*_10_) to quantify the relative evidence the data provide in favor of the alternative hypothesis (*H*_1_) over the null (*H*_0_), which we interpret following Jeffreys (1961). All tests are non-directional as defined by a symmetric prior (unless otherwise indicated).

### Correlations

For testing linear correlations with Pearson’s *r*, the Bayesian test is based on Jeffrey’s (1961) test for linear correlation and assumes a shifted, scaled beta prior distribution 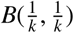 for *r*, where the scale parameter is set to 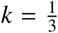 (Ly, Verhagen, & Wagenmakers, 2016).

### Bayesian model selection

We use a Bayesian model selection framework designed for group studies (Rigoux et al., 2014; Stephan et al., 2009) to evaluate our models. Intuitively, it can be described as a random-effect analysis, where models are treated as random effects and are allowed to differ between subjects. Assuming that there is a fixed but unknown distribution of models in the population, the goal is to determine the probability of each model being more frequent in the population than all other models in consideration. This is modelled hierarchically, using variational Bayes to estimate the parameters of a Dirichlet distribution describing the posterior probabilities of each model *P*(*m*_*k*_ |**y**) given the model evidence **y**, which is approximated using the cumulative out-of-sample negative log likelihoods of each participant, obtained from cross-validated maximum likelihood estimation (Fong & Holmes, 2020). The exceedance probability (*xp*) is thus defined as the posterior probability that the frequency of a given model 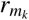 is larger than all other models 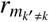 under consideration:

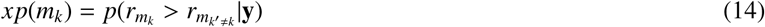

Rigoux et al. (2014) extends this approach by correcting for chance, based on the Bayesian Omnibus Risk (*BOR*), which is the posterior probability that all model frequencies are equal. This produces the *protected exceedance probability* (*pxp*) reported throughout this article.

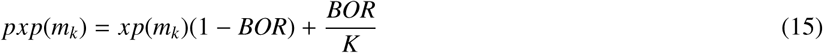

### Regression

All Bayesian mixed effects regression models were implemented in brms with generic weakly informative priors (Bürkner, 2017) using No-U-Turn sampling (Hoffman & Gelman, 2014) with the proposal acceptance probability set to .99. In all cases, participant id was used as a random intercept. All fixed effects were also entered as random slopes following a maximal random structure approach (Barr, Levy, Scheepers, & Tily, 2013). This allows us to compute a Bayes factor comparing each model against a null model, which used the same random effect structure but with the target fixed effect omitted. The Bayes factor was computed using bridge sampling (Gronau, Singmann, & Wagenmakers, 2017) as a method to approximate the marginal likelihood of both models. All models were estimated over four chains of 4000 iterations, with a burn-in period of 1000 samples.

**Table A1.** Experiment 1: Mixed Effects Models

|  | Judgment Accuracy | Judgment Accuracy | Confidence Rating | Confidence Rating |
| --- | --- | --- | --- | --- |
|  | *Est.* | *Est.* | *Est.* | *Est.* |
| Intercept | 12.77<br>[11.73, 12.80] | 14.89<br>[13.50, 16.31] | 6.62<br>[6.14, 7.11] | 8.26<br>[7.95, 8.58] |
| Observed Nodes | -0.60<br>[-0.79, -0.41] |  | 0.23<br>[0.17, 0.30] |  |
| Confidence Rating |  | -0.66<br>[-0.83, -0.49] |  |  |
| GP Uncertainty |  |  |  | -1.80<br>[-2.50, -1.12] |
| **Random Effects** |  |  |  |  |
| $\sigma^2$ | 4.86 | 4.77 | 2.61 | 2.40 |
| $\tau_{00}$ | 73.37 | 73.52 | 3.37 | 3.58 |
| ICC | 0.06 | 0.06 | 0.44 | 0.40 |
| N | 100 | 100 | 100 | 100 |
| Observations | 3000 | 3000 | 3000 | 3000 |
| Bayesian $R^2$ | .07 | 0.09 | 0.46 | 0.43 |

*Note:* We report the posterior mean and 95% highest posterior density (HPD) interval below in brackets. $\sigma^2$ indicates the individual-level variance, $\tau_{00}$ indicates the variation between individual intercepts and the average intercept, and ICC is the intraclass correlation coefficient.

**Table A2.** Experiment 2: Mixed Effects Models

|  | P(repeat)<br>*Odds Ratio* | Graph Distance<br>*Est.* | Reward<br>*Est.* | Eigen Centrality<br>*Est.* |
| --- | --- | --- | --- | --- |
| Intercept | 0.00<br>[0.00,0.00] | 10.59<br>[9.85, 11.36] | 78.61<br>[76.65, 80.51] | 0.15<br>[0.14, 0.17] |
| Previous Reward | 1.13<br>[1.12,1.14] | -0.11<br>[-0.12, -0.10 ] |  |  |
| Trial |  |  |  | -0.003<br>[-0.003, -0.002] |
| Eigen Centrality |  |  | -26.65<br>[-31.17, -22.04] |  |
| **Random Effects** |  |  |  |  |
| $\sigma^2$ | -0.01 | 1.37 | 65.15 | 0.003 |
| $\tau_{00}$ | 0.25 | 18.47 | 496.55 | 0.04 |
| ICC | -0.02 | 0.07 | 0.12 | 0.07 |
| N | 98 | 98 | 98 | 98 |
| Observations | 22050 | 22050 | 22050 | 22932 |
| Bayesian $R^2$ | .58 | .42 | .17 | .08 |

*Note:* We report the posterior mean and 95% highest posterior density (HPD) interval below in brackets, except the first column, which is a logistic regression reporting the odds ratio and the respective 95% CI. $\sigma^2$ indicates the individual-level variance, $\tau_{00}$ indicates the variation between individual intercepts and the average intercept, and ICC is the intraclass correlation coefficient.

**Table A3.** Experiment 2: Bonus Round

|  | Judgment<br>*Est.* | Judgment<br>*Est.* | Judgment<br>*Est.* | Confidence<br>*Est.* |
| --- | --- | --- | --- | --- |
| Intercept | -12.38<br>[-26.16,-0.89] | 5.15<br>[-1.04, 11.11] | 5.24<br>[-2.37, 12.33] | 7.72<br>[7.23,8.22] |
| GP Prediction | 1.16<br>[0.93,1.43] |  |  |  |
| GP Uncertainty |  |  |  | -3.13<br>[-4.37, -1.92] |
| dNN Prediction |  | 0.80<br>[0.69,0.92] |  |  |
| kNN Prediction |  |  | 0.79<br>[0.66,0.94] |  |
| **Random Effects** |  |  |  |  |
| $\sigma^2$ | 51.14 | 53.38 | 63.86 | 2.55 |
| $\tau_{00}$ | 275.06 | 274.38 | 272.84 | 4.37 |
| ICC | 0.16 | 0.16 | 0.17 | 0.37 |
| N | 98 | 98 | 98 | 98 |
| Observations | 980 | 980 | 980 | 980 |
| Bayesian $R^2$ | .32 | .33 | .33 | .50 |

*Note:* Note that the Bayesian mean tracker (BMT) is not included, since it invariably makes the same prediction for all unobserved options based on the prior mean and prior variance. The bonus round only consisted of judgments about unobserved options. GP, Gaussian process; dNN, *d*-Nearest Neighbors; kNN, *k*-Nearest Neighbors. We report the posterior mean and 95% highest posterior density (HPD) interval below in brackets. $\sigma^2$ indicates the individual-level variance, $\tau_{00}$ indicates the variation between individual intercepts and the average intercept, and ICC is the intraclass correlation coefficient.

## Appendix B Experiment 1 model supplement

### Predictive performance

The predictive performance is defined in terms of log likelihood of the out-of-sample predictions, which is a monotonic transformation of the mean squared prediction error by assuming a Gaussian probability density:

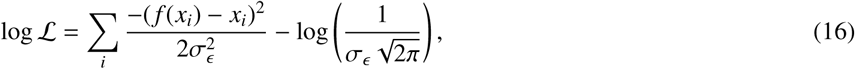

where each *x*_*i*_ are the participant judgments, each *f* (*x*_*i*_) are the out-of-sample model predictions, and we set σ*ϵ* = 1.

### Correspondence between participant judgments and model predictions

In addition to comparing prediction accuracy (Fig. 3d-e), we also examined the correspondence between participant judgments, the true underlying values, and model predictions. Figure B1a shows a scatter plot comparing participant judgments to the true target value (*r* = .59, *p* < .001, *BF*_10_ > 10^9^), which shows participants were well calibrated to ground truth. Figure B1b-d provides similar scatter plots showing the correspondence between each model’s predictions and participant judgments. Overall, the GP had the highest correlation with participant judgments (*r* = .71, *p* < .001, *BF*_10_ > 10^9^), followed by the dNN (*r* = .68, *p* < .001, *BF*_10_ > 10^9^) and kNN models (*r* = .67, *p* < .001, *BF*_10_ > 10^9^). Comparing *z*-transformed correlation coefficients computed at the individual level, we find that the GP predictions were more correlated to participant judgments than the kNN (*t*(99) = 8.6, *p* < .001, *d* = 0.9, *BF*_10_ > 10^9^), but with no differences between the GP and dNN (*t*(99) = −1.0, *p* = .335, *d* = 0.0, *BF* = .17).

**Figure B1.**
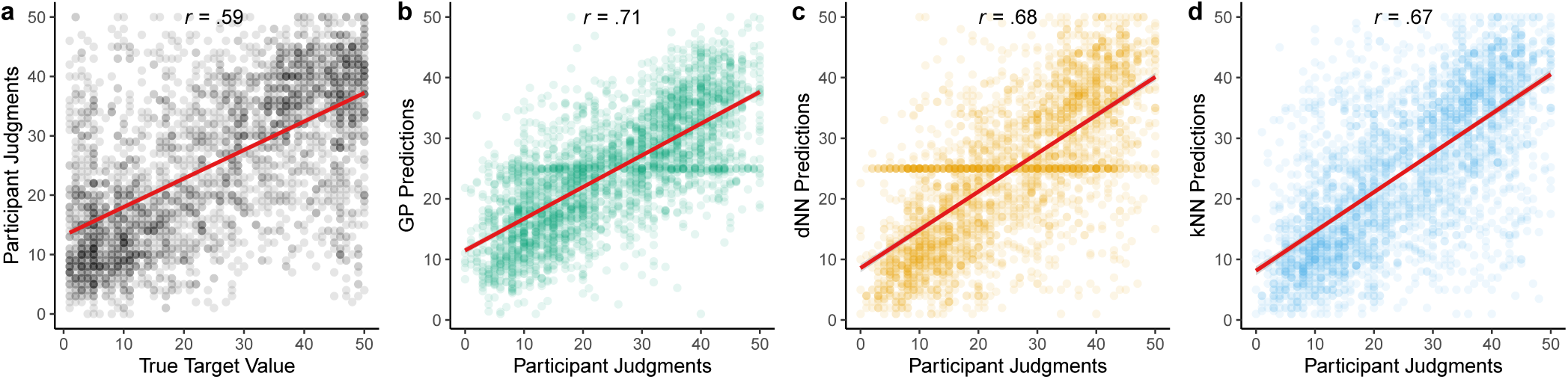
Experiment 1 correspondence between participant and model predictions. **a**) Participant judgments against the ground truth. **b-d**) Model predictions compared against participant judgments. Each dot is a single judgment and the red line is a linear regression. The Pearson correlation coefficient is shown above (*BF*_10_ > 10^9^ in all cases). GP, Gaussian process; dNN, *d*-Nearest Neighbors; kNN, *k*-Nearest Neighbors

## Appendix C Eigen Centrality

We analyzed search behavior from Experiment 2 as a function of the connectivity of the sample nodes, which we quantify using eigen centrality (EC; Bonacich, 1972). Intuitively, EC quantifies the connectivity of a node similar to how Google’s PageRank (Langville & Meyer, 2011) quantifies webpages based on the number and quality of hyperlinks. Nodes with higher EC are those that exert higher influence on the network, by being connected to other nodes that are themselves highly central in the network. The ECs of each node in a graph *x*_*i*_ ∈ **x** are defined by the normalized eigenvector belonging to the largest eigenvalue λ of the adjacency matrix *A*, fulfilling the identity:

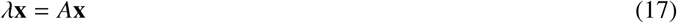

Compared to the overall distribution of ECs in the task, participants systematically selected nodes with lower EC (one-sample *t*-test: *t*(97) = −9.0, *p* < .001, *d* = 0.9, *BF* = 3.2 × 10^11^; Fig. C1a). Participants also increasingly selected lower EC nodes over successive trials (Bayesian mixed model: *b*_*trial*_ = −0.003, 95% HPD: [−0.003, −0.002], *BF*_10_ = 1.6 × 10^11^; Table A2; Fig. C1b). While EC was not predictive of expected rewards in the underlying task environment (*r* = − .03, *p* = .109, *BF* = .17; Fig. C1c), we found a systematic relationship in the choices made by participants: lower EC nodes sampled by participants had higher reward values (*b*_*eigenCentrality*_ = −26.54, 95% HPD: [−31.17, −22.04], *BF*_10_ = 1.9 × 10^20^; Table A2; Fig. C1d). This is perhaps because nodes with lower EC tended to have more eccentric reward values. Indeed, the highest and lowest rewards across environments had very similar average EC values, of 0.06 and 0.05, respectively. Thus, this trend of preferentially sampling less central nodes may reflect a high-risk high-reward heuristic (Leuker, Pachur, Hertwig, & Pleskac, 2018), which combined with generalization proved to be an adaptive search strategy.

**Figure C1.**
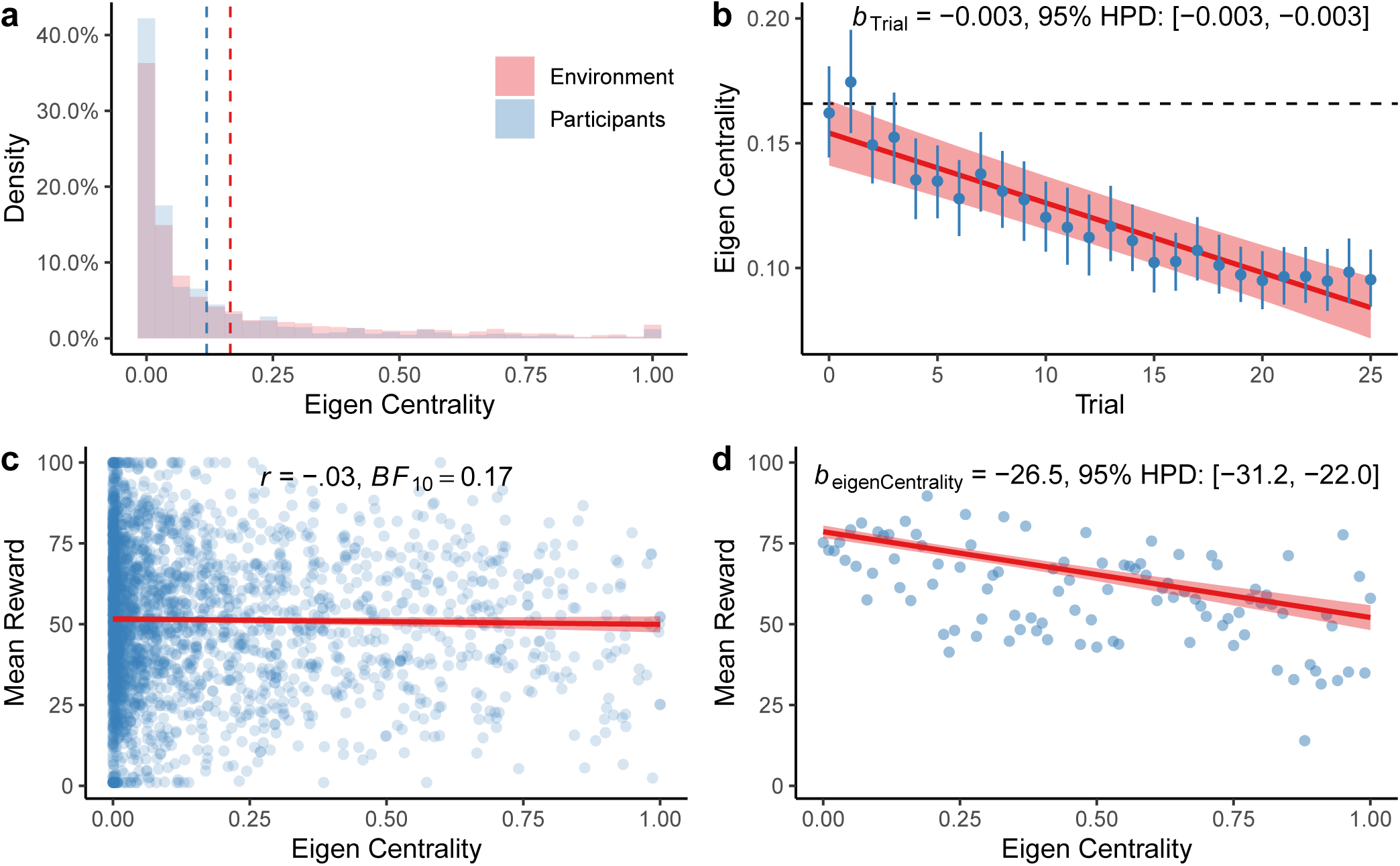
Eigen Centrality. **a**) Distribution of EC values of selected options (blue) compared to the ground truth of the task (red). The vertical dashed lines indicate the means of each distribution. **b**) Participants preferentially selected nodes with lower EC over subsequent trials. Each dot is the aggregate mean (±95% CI) and the red line shows the group-level effect of a Bayesian mixed model (Table A2), with the ribbon showing the 95% CI. The dashed horizontal line indicates the mean eigen centrality across all nodes in the task. **c**) EC was not predictive of rewards in the task. **d**) However, from the nodes sampled by participants, those with lower EC corresponded to higher rewards. Each dot is dot is the aggregate mean (calculated at intervals of 0.01) and the red line is the group-level effect of a Bayesian mixed model (Table A2), with the ribbon indicating the 95% CI.

## Appendix D Experiment 2 model supplement

Figure D1 provides an overview of parameter estimates for each model, while Figure D2 shows how different parameter estimates were related to different levels of predictive accuracy. In addition, Figure D3 compares the difference in out-of-sample prediction error between the GP and each other model as a function of performance on the bandit task.

### Parameter estimates

#### GP

The GP used the diffusion parameter (α) to define the extent of generalization, where larger values of α implied a wider influence of observed rewards over the graph structure. While the superior predictive accuracy of the GP over alternative models provided evidence for generalization, α estimates were systematically lower than the underlying value of α = 2 used to generate the environments (*t*(97) = −13.5, *p* < .001, *d* = 1.4, *BF*_10_ = 9.3 × 10^20^). Thus, undergeneralization rather than overgeneralization was the norm, consistent with previous findings of a beneficial a bias towards undergeneralization in a similar search context (Wu, Schulz, Speekenbrink, et al., 2018). Additionally, the exploration bonus (*β*) estimated how participants traded off between exploring uncertain options vs. exploiting options with high expectations of reward. We found that the estimated *β* values were substantially larger than the lower bound (*t*(97) = 4.5, *p* < .001, *d* = 0.5, *BF*_10_ = 949), providing further evidence for directed exploration. Lastly, we also define a stickiness parameter (ω), which captures an aspect of the high rates of repeat clicks by adding an additional bonus to the value of the last selected options.

#### BMT

In comparison, while the BMT also made uncertainty estimates, these were defaulted to the prior variance (*v*_0_ = 500) for all unobserved options. Thus, the BMT made the predictions uncertainty predictions for nodes near and far from previous observations. In contrast to the GP model, we found little evidence for directed exploration using the BMT, with *β* estimates only marginally different from the lower bound of .007 (*t*(97) = 2.1, *p* = .038, *d* = 0.2, *BF*_10_ = .92). The BMT also made similar use of the stickiness parameter compared to the GP (*t*(97) = 0.7, *p* = .460, *d* = 0.1, *BF*_10_ = .15).

#### Nearest-neighbors

The dNN generated predictions by averaging the rewards of observed nodes within a distance of *d*. The mean estimate of distance was *d* = 2.4, although the mode and median were both 1. Thus, the dNN predominately made predictions solely based on observations of directly connected nodes. Nonetheless, it was still able to predict participant choices fairly accurately. While the dNN had no access to directed exploration, we nevertheless find similar levels of random exploration (τ: *t*(97) = 1.2, *p* = .230, *d* = 0.1, *BF*_10_ = .23) and stickiness (ω: *t*(97) = −1.0, *p* = .317, *d* = 0.1, *BF*_10_ = .18) compared to the GP. Thus, one potential source of the gap in simulated learning performance compared to the GP (Fig. 5e), could be due to the dNN lacking a form of directed exploration. The kNN model also performed similar to the dNN, by averaging the *k* nearest nodes rather than selecting nodes at a fixed distance. The mean number of neighbors was *k* = 3, and with a mode and median of 2. Thus, like the dNN, generalizations were on the basis of integrating a small number of other observations. The dNN and kNN also shared similar levels of both undirected exploration (*t*(97) = 1.8, *p* = .077, *d* = 0.3, *BF*_10_ = .52), and stickiness (*t*(97) = 1.1, *p* = .290, *d* = 0.2, *BF*_10_ = .19).

**Figure D1.**
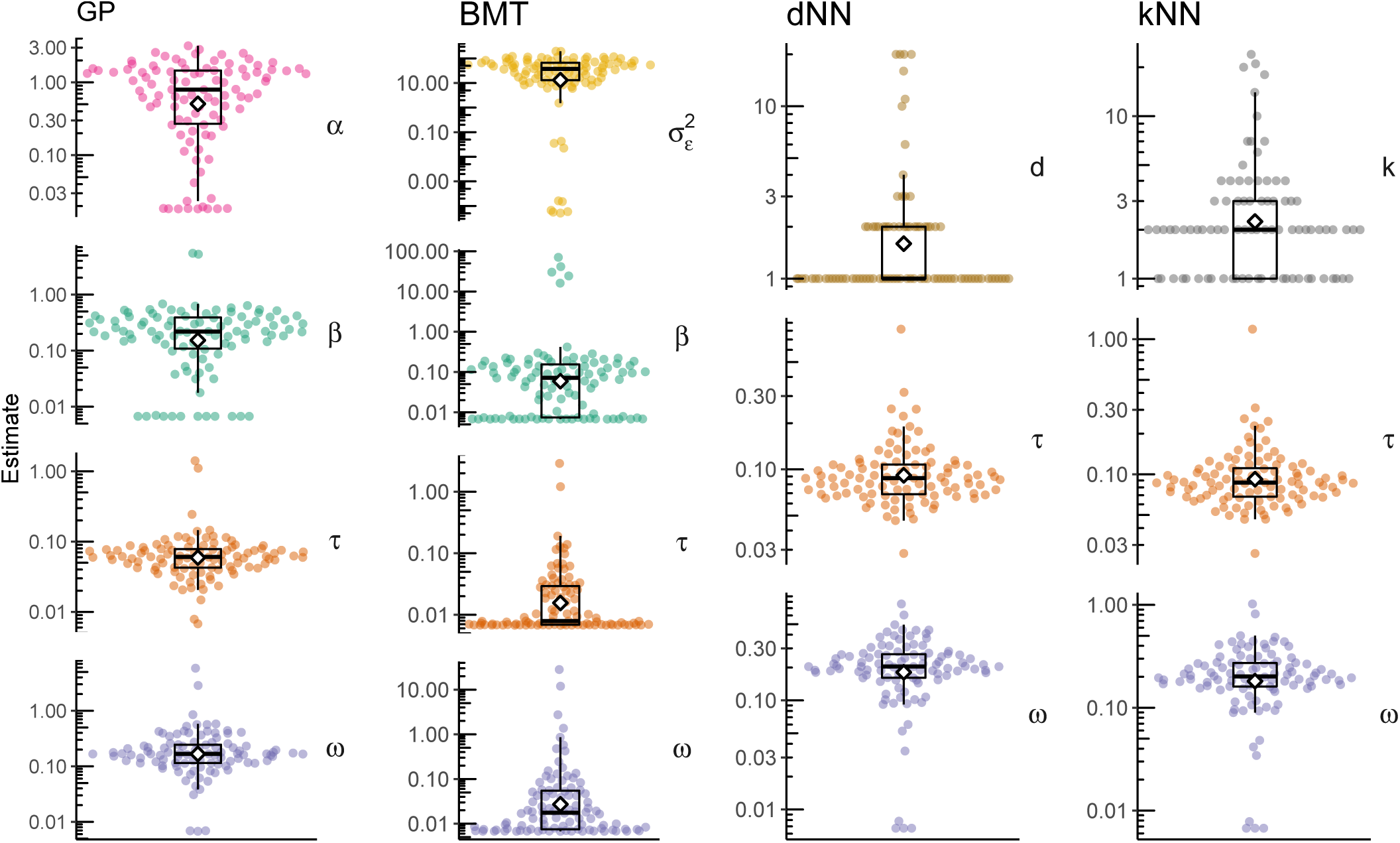
Experiment 2 parameter estimates. Each dot is the median cross-validated estimate for a single participant. The diamond indicates group mean and Tukey box plots show the median and 1.5 inter-quartile range. Note that values of *d* and *k* are restricted to the set of natural numbers. GP, Gaussian process; BMT, Bayesian mean tracker; dNN, *d*-Nearest Neighbors; kNN, *k*-Nearest Neighbors.

**Figure D2.**
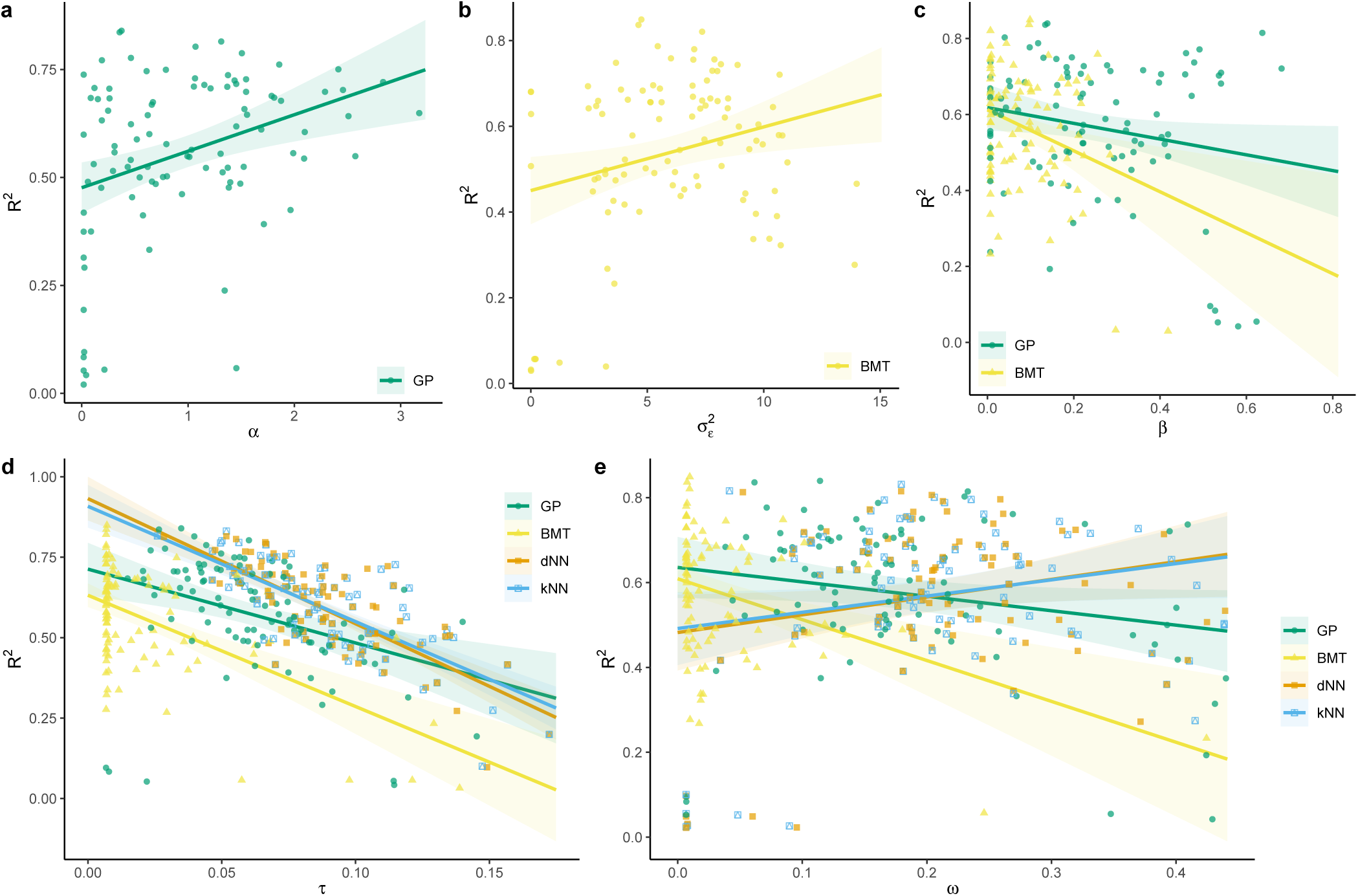
Experiment 2 model parameters and predictive accuracy. Each panel compares median per-participant parameter estimates against predictive accuracy (*R*^2^), as an intuitive measure of objective model performance. Predictive accuracy compares the cumulative out-of-sample negative log likelihood of any given model *k* against a random model: *R*^2^ = 1 − log *ℒ*_*k*_/ log *ℒ*_*rand*_. Intuitively, *R*^2^ = 0 indicates a model is equivalent to random chance, while *R*^2^ = 1 is a theoretically perfect model. Each dot is a single participant, with linear regression lines added (ribbon indicates standard error). **a**) The diffusion parameter α measuring the level of generalization, where higher levels of generalization corresponded to better model predictions (*r* = .34, *p* < .001, *BF*_10_ = 54) **b**) The inverse sensitivity parameter 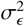, where higher estimates (i.e., smaller learning updates) corresponded to better model predictions (*r* = .25, *p* = .012, *BF*_10_ = 4.7). **c**) The exploration bonus *β* controls the level of *directed* exploration. For both GP and BMT models, higher levels of directed exploration corresponded to worse model predictions (*BF*_10_ > 100). **d**) The softmax temperature parameter τ controls the level of *random* exploration. For all models, temperatures corresponded to worse model predictions (*BF*_10_ > 100). **e** The stickiness parameter ω added a bonus to the previously selected option, making it more likely to choose the same option on the next trial. For the GP and BMT models, stickiness was correlated with worse model model predictions (*BF*_10_ > 100), whereas there was no relationship for the dNN and kNN models (*BF*_10_ < 1). GP, Gaussian process; BMT, Bayesian mean tracker; dNN, *d*-Nearest Neighbors; kNN, *k*-Nearest Neighbors.

**Figure D3.**
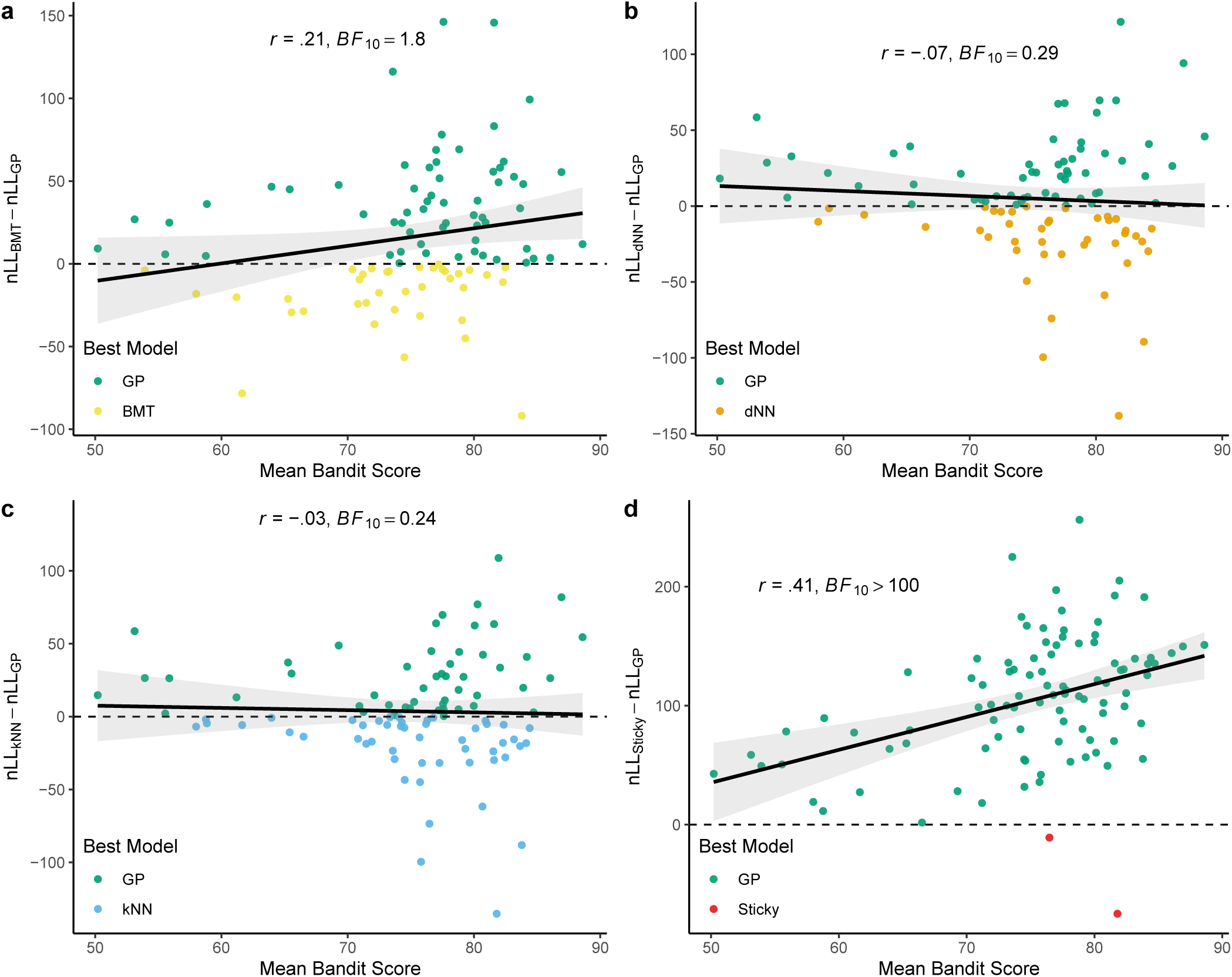
Experiment 2 Model Differences and Score. Each panel compares the difference in out-of-sample prediction error, measured as negative log likelihood (nLL), of two models as a function of mean performance on the bandit task. Each dot is a single participant, with the Pearson correlation shown above and a linear regression line added to the plot (ribbon indicates standard error). The color of each dot indicates the model with the lower nLL. **a**) GP vs. BMT. **b**) GP vs. dNN. **c**) GP vs. kNN. **d**) GP vs. Stickiness and softmax model. GP, Gaussian process; BMT, Bayesian mean tracker; dNN, *d*-Nearest Neighbors; kNN, *k*-Nearest Neighbors.

## Footnotes

1 We follow the convention of setting the mean function to zero, such that the GP prior is fully defined by the kernel

2 In practice, the matrix exponentiation in Eq. 5 can be computed by first decomposing **L** into its eigenvectors {*u*_*i*_} and eigenvalues {λ_*i*_}, and then substituting matrix exponentiation with real exponentiation using 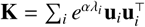.

3 Previous results reported in Wu, Schulz, and Gershman (2019) suffered from numerical instability during matrix exponentiation when computing the diffusion kernel, and thus yielded different estimates.

